# Confounding Fuels Misinterpretation in Human Genetics

**DOI:** 10.1101/2023.11.01.565061

**Authors:** John W. Benning, Jedidiah Carlson, Olivia S. Smith, Ruth G. Shaw, Arbel Harpak

## Abstract

The scientific literature has seen a resurgence of interest in genetic influences on human behavior and socioeconomic outcomes. Such studies face the central difficulty of distinguishing possible causal influences, in particular genetic and non-genetic ones. When confounding between possible influences is not rigorously addressed, it invites over- and misinterpretation of data. We illustrate the breadth of this problem through a discussion of the literature and a reanalysis of two examples. Clark (2023) suggested that patterns of similarity in social status between relatives indicate that social status is largely determined by one’s DNA. We show that the paper’s conclusions are based on the conflation of genetic and non-genetic transmission (for example, of wealth) within families. Song & Zhang (2024) posited that genetic variants underlying bisexual behavior are maintained in the population because they also affect risk-taking behavior, thereby conferring an evolutionary fitness advantage through increased sexual promiscuity. In this case, too, we show that possible explanations cannot be distinguished, but only one is chosen and presented as a conclusion. We discuss how issues of confounding apply more broadly to studies that claim to establish genetic underpinnings to human behavior and societal outcomes.

## Introduction

People vary remarkably in behavior and social outcomes. This variation sparks curiosity about its causes, and for the past 150 years, scholars have debated the extent to which it arises due to underlying genetic differences. In the 19th century, Galton (*1*) found strong resemblance between parents and their offspring in measures of social status and on that basis inferred that genetics is the most likely root cause, a school of thought described broadly as “hereditarianism” (see *2*). As is now well appreciated, Galton’s inference neglected the fact that parents transmit not only genetic material to their offspring, but also wealth, place of residence, knowledge, religion, culture, and more. For such attributes, transmission within families can parallel genetic transmission (**Fig. 1a**) (*3–20*), often leading genetic and non-genetic transmission to be indistinguishable in observational data. A long history of scholarship has highlighted this type of confounding and how it impedes inference of the causes of phenotypic variation (*21–28*). Studies without molecular genetic data are particularly susceptible to confounding, because they offer little to no signal that could be argued to reflect only genetic or only non-genetic transmission (*21, 22, 25, 27, 29*).

**Figure 1.**
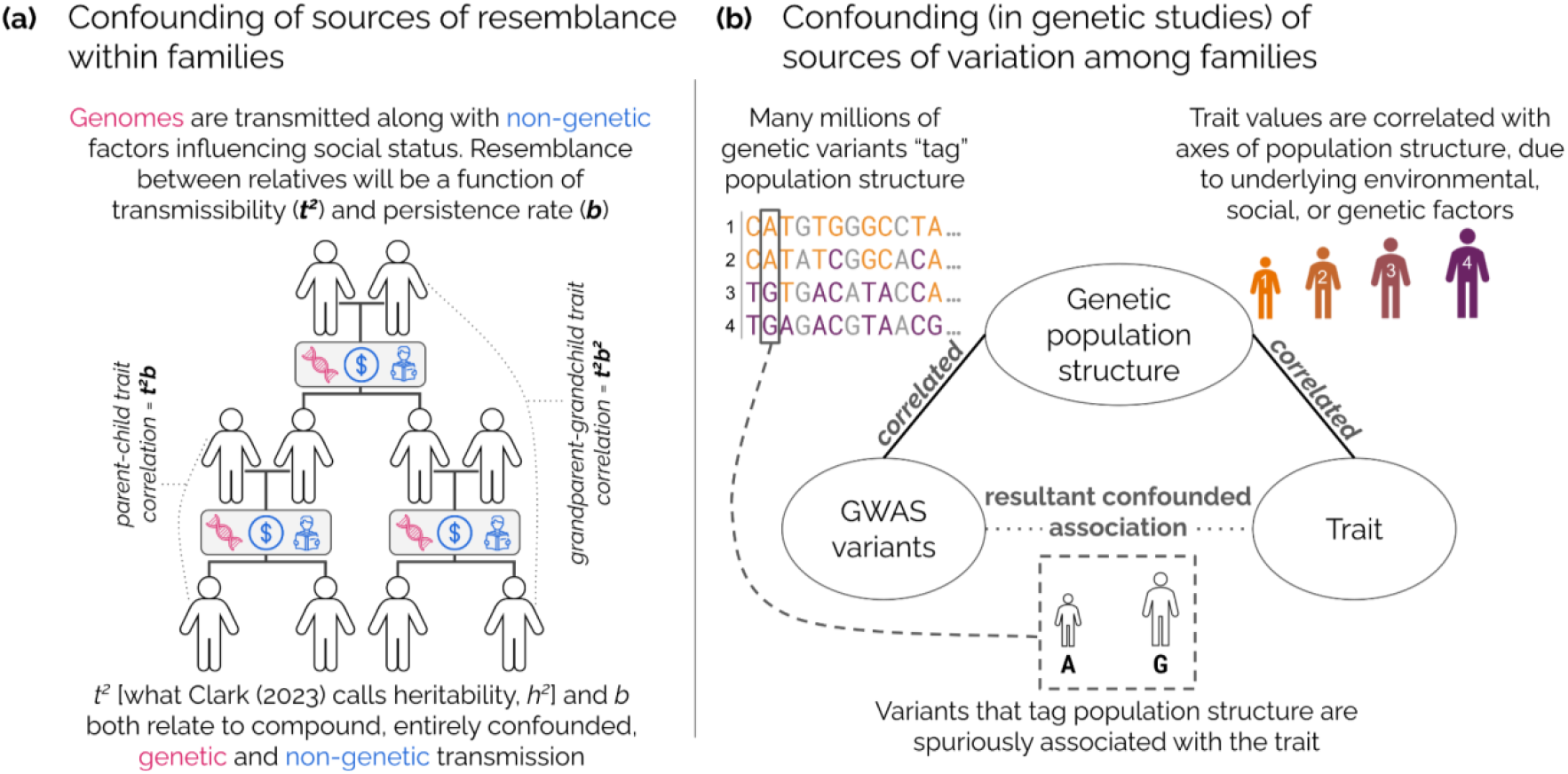
Confounding between genetic and non-genetic factors influencing traits. (a) Confounding within families. Non-genetic transmission can parallel genetic transmission and their respective effects are confounded in observational data. Illustrated is a model where a trait value is the sum of an inherited component from parents and random noise. Under this model, the expected resemblance between relatives depends on transmissibility (*t*^*2*^, the portion of trait variation attributable to the transmitted component) and a rate of decay across genealogical distance (the “persistence rate,” *b*, which increases with increasing degree of assortative mating). Ignoring the confounding of genetic and non-genetic transmission in the data, Clark (2023) misassigns all transmission as genetic heritability and all assortative mating to be on a latent “social genotype”. **(b) Confounding among families induces biases in GWAS**. “Population structure confounding” in genomic data relates to correlations between the structure of genetic relatedness in a GWAS sample (exemplified by the orange-to-purple gradient) and the phenotype studied. Here we show genetic sequences from individuals 1-4 at top left, with their attendant phenotypes (height) at top right. For a given genetic variant, individuals with purple alleles will tend to be taller than those with orange alleles, regardless of the variant’s causal effect on height. This confounding affects any variants that reflect this axis of genetic population structure— typically many millions of variants. While researchers often use methods that adjust for population structure in an attempt to avoid spurious associations, the extent of residual confounding in GWAS remains unclear.

This manuscript is motivated by the fact that confounding is still frequently overlooked or downplayed in reports about genetic causes of human behavior and socioeconomic outcomes. We demonstrate instances of confounding that underlie misinterpretation in current, high-profile primary literature that have been drawing media attention (ranking in the 99th percentile of Altmetric Attention Scores for papers of a similar age) through reanalysis of their data. We begin with a recent publication that made claims about genetic determinism of social status (*30*) in the absence of molecular genetic data or viable strategies for disentangling genetic from non-genetic contribution (**Box 1i**). We then discuss sources of confounding undermining causal inference based on genome wide association studies (GWAS) for behavior and social outcomes, with a focus on confounding via population stratification (**Box 1ii**,**iii**). Lastly, we consider the impacts of related errors in causal inference stemming from data preparation and other analysis choices (“analytical confounding”; **Box 1iv**). We illustrate these problems using a recent study (*31*) that purported to explain the evolutionary maintenance of genetic variation affecting bisexual behavior.

#### Box 1: What is confounding?

Consider the hypothesis that a change in variable A causes a change in variable B, denoted A → B. Another variable, C, that affects both A and B, is called a confounder: it renders the effect of A on B indistinguishable from the effect of C on B. In the presence of confounding, one cannot reliably estimate causal influences (*32, 33*).

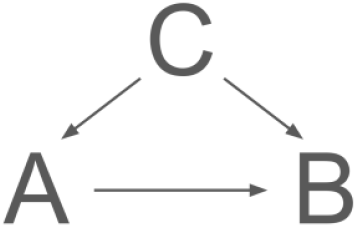

#### Confounding data

Sometimes, variables that are not confounders in the population can still act as confounders in a dataset.

This can happen when the structure of the data prevents variation in one variable while holding another constant—so even if C does not cause A in the population, in the dataset, one cannot vary A independently of C. Confounding can also arise when multiple causal factors (e.g., A_1_ and A_2_) are aggregated into a single variable A. If only A is measured, and A_1_ is interpreted as the causal driver, associations arising from A_2_ may be mistakenly attributed to A_1_ (this is one mode of what is sometimes termed “proxy confounding” (*34, 35*)). In both cases, the structure of the data—rather than solely the causal mechanisms that exist in the population—determines what is confounded.

#### Modes of confounding discussed in this work

| Case discussed in this manuscript | (A) Putative cause | (B) Outcome | (C) Confounder |
| --- | --- | --- | --- |
| i. Familial transmission (Clark 2023; **Fig. 1a**) | Genetic variants | Social status | Non-genetic factors transmitted within families, such as wealth, community, and culture.<br>These influences covary with genetic variants across individuals, forming a single, inseparable 'transmissible component'. Thus, this is a form of proxy confounding: even though the non-genetic factors do not influence genotypes in the population, the two are bundled together in the same variable (individual identifier) in the data. |
| ii. Genetic confounding (**Fig. 1b**) in a GWAS for height | Focal genetic variant | Height | Other genetic variants influencing height that are correlated (in “linkage disequilibrium”) with the focal variant. |
| iii. Environmental/social confounding (**Fig. 1b**) in a GWAS for asthma | Focal genetic variant | Asthma | Environmental or social influences on asthma, such as exposure to pollutants, that correlate with the focal variant |
| iv. Analytical confounding via choice of hypotheses to consider (Song and Zhang 2024; **Fig. 4a,b**) | Effects of genetic variants, associated with bisexual behavior, on risk-taking behavior | Effects of these genetic variants on number of children (as a proxy for evolutionary fitness) | Effects of these variants on traits and outcomes other than risk taking (none were considered in the study) |

#### In observational studies, confounding may be insurmountable

Pearl’s backdoor criterion (*34, 36*) provides a formal way to estimate the causal effect of A on B in observational data: we must block all “backdoor paths”—paths that connect A and B through common causes rather than through the effect of A on B—by conditioning on common causes of both A and B, while avoiding conditioning on variables affected by A.

Applying this criterion can be difficult or impossible in observational studies. One reason is that we may not observe, or know of, the relevant confounding variables. Even when we do, these variables may be, in addition to affecting A, affected by A themselves, meaning that we cannot apply Pearl’s backdoor criterion.

### Confounding fuels hereditarian fallacies

A recent publication (*30*) analyzed familial correlations in a dataset of socioeconomic measures (e.g., occupational status, house value, literacy) from a selection of records spanning the 18th to 21st centuries in England. In it, a quantitative genetic model is fitted to these observed correlations [(*37, 38*); **Supplementary Note 1**]. Based on this fit, (*30*) infers that social status persists intergenerationally because of strong assortative mating on a status-determining genotype (or “social genotype” as the author has termed it in previous work (*39*)). Further, the paper argues that the persistence of social status within families—and persistence of differences in status among families—have been largely unaffected by changes in social policy in the last four centuries. In a subsequent commentary about this work (*40*), the author presents the results of as providing strong support for a hereditarian interpretation. In doing so, he appeals to the metaphor of a “genetic lottery” underlying social outcomes.

Here, we discuss the failure to account for the confounding of genetic and non-genetic transmission (**Fig. 1a**) that, together with other core flaws of the analysis (**Fig. 2a-b**), fuels the hereditarian claims in (*30*) (see our discussion of other misinterpretations, errors, and incongruencies in (*30*) in **Supplementary Notes 2-7**; **Tables S1-S3**; **Figs. S1-S13**). We also demonstrate that familial status correlations varied substantially over the time period examined, generally decreasing (**Fig. 2c**). This finding contrasts with conclusions in (*30*), based on the same data, that social mobility has been stagnant. As we show below, the analyses in (*30*) do not establish the contribution of genetics to social status.

**Figure 2.**
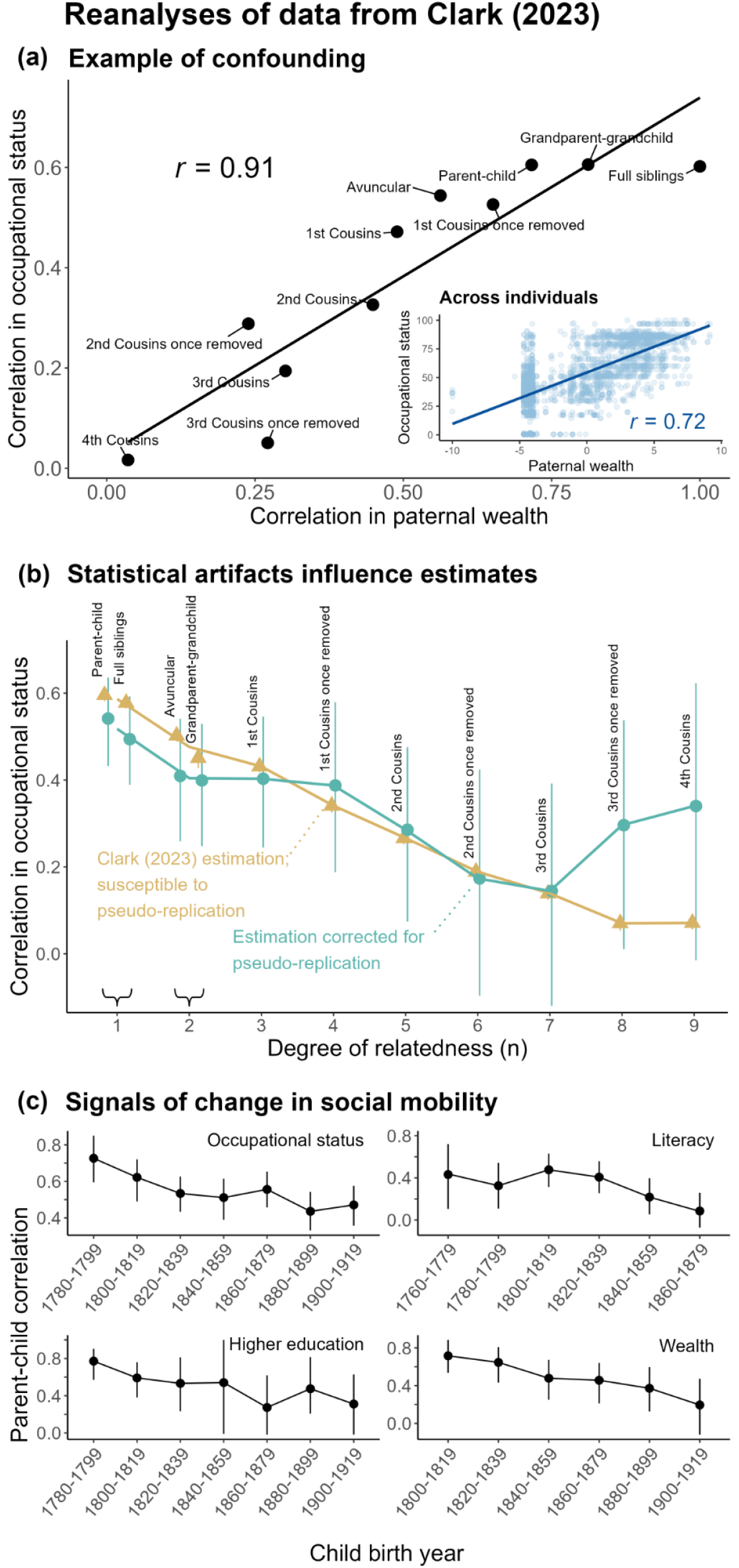
Reanalyses of data from Clark (2023) challenge the paper’s claims. (a) Example of confounding between genetic and non-genetic transmission. The relationship between paternal wealth and a measure of social status suggest at least one source of confounding between genetic and non-genetic transmission. Across relative pairs, correlation in occupational status is highly correlated (Pearson’s *r* = 0.91) with those relatives’ correlation in paternal wealth. Inset shows that individual occupational status is strongly correlated (Pearson’s *r* = 0.72) with father’s wealth. Excluding outliers in paternal wealth (1.5×IQR rule) had only a small impact (a change of less than 0.01) on the Pearson correlations reported here. Plots show data for individuals born 1780-1859 and their fathers. For the cohort of individuals born after 1859, the correlations change to *r* = 0.72 for relative pairs and *r* = 0.60 for individuals. (*30*) estimated wealth from probate records. The log of estimated wealth was mean-centered with respect to 5-year bin means. Individuals not probated due to insufficient wealth were assigned a value of half the minimum probate requirement for the time period.) **(b) Pseudoreplication distorted estimates of familial correlations**. Familial correlations (95% CI) in occupational status (1780-1859) using the approach employed by (*30*) (in gold) involved pervasive, non-uniform pseudoreplication (Supplementary Note 6). For example, the (1780-1859) occupational status correlation for fourth cousins is calculated from 17,382 pairs, derived from only 1,878 unique individuals. In teal we show conservative estimates using only a single relative pair per surname (means and 95% CI over 1000 bootstrap samples are plotted for each familial correlation), which are therefore not susceptible to pseudoreplication. Distant cousins show dramatically higher correlations after adjusting for pseudoreplication. We note that additional biases may exist that are not addressed with this adjustment (Supplementary Note 6) **(c) Signals of change in social mobility**. Parent-offspring correlations in multiple status measures generally decrease over time in (*30*)’s data, in contrast to claims of stagnant social mobility made in the original paper. To mitigate pseudoreplication, we calculated correlations using one pair from each surname [as in (b)]. Shown are average correlations (95% CI) across 500 bootstrap iterations of correlation estimation. Fig. S13 shows two complementary analyses estimating correlations either without accounting for pseudoreplication, or using percentile ranks—both result in similar trends.

#### Confounding between genetic and non-genetic transmission

Inferences in (*30*) are based on a linear regression model derived from quantitative-genetic theory developed by R.A. Fisher (*37, 38*) (**Supplementary Note 1**) and the model

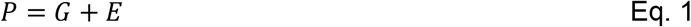

where an individual’s phenotype, *P*, is the sum of separable genotypic (*G*) and environmental (*E*) influences on it. Because genotypes are transmitted from parents to offspring, genetic parameters can be inferred from correlations between relatives, on the condition that environmental influences are independent and random with respect to genotypes. Fisher (1918) formally showed that under this model, the expected correlation in a trait between pairs of individuals of a defined relationship is a function of the genealogical relationship between the relatives, the trait’s heritability (*h*^*2*^), and the extent of assortative mating in the population (*b*). (*h*^*2*^ is the fraction of phenotypic variance due to additive genetic variance, commonly referred to as “narrow-sense” heritability.)

Crucially, to interpret the model parameters *h*^*2*^ and *b* as relating to genetic effects, Fisher’s model assumes that there are no non-genetic (material, environmental, or cultural) influences on a trait that are systematically shared or transmitted between relatives. This assumption is valid in an experimental setting, for instance, in which genotypes are randomized with regard to environment.

However, in humans that assumption is nonsensical. Non-genetic transmission is ubiquitous for social and behavioral traits. Traits may be transmitted directly between relatives (e.g., literate parents teaching their children how to read) (*5*), or via indirect mechanisms such as “ecological inheritance,” where the trait value of an offspring is influenced by the environmental conditions bestowed by their parents (e.g., familial wealth influencing educational opportunities) (*8, 41*). When genotypes cannot be randomized over environments, true genetic effects are much more difficult to separate from other factors underlying phenotypic resemblance between relatives (*22*). In (*30*), for instance, the assumption of no systematic non-genetic transmission implies that similarity in house value among relatives (one of the measures of social status analyzed) is solely due to shared genes, and does not arise from similarity in parental wealth, the inheritance of wealth or property, or having learned from one’s relatives about investment.

In fact, we found evidence of strong confounding between genetic and non-genetic contributions to familial resemblance in the data used in (*30*). The paper acknowledges the inheritance of material wealth from one’s parents as an example of non-genetic transmission only when treating wealth itself as the focal status measure. For other measures studied, the effect of familial wealth on social status is ignored. Yet familial wealth can obviously influence a wide range of conditions that affect offspring (e.g., healthcare, place of residence, access to tutors, social circles, etc.) (*42*– *46*). Consistent with this intuition, we found that all seven status measures analyzed in (*30*) are substantially correlated with an individual’s father’s wealth (Pearson *r* ranging from 0.19 - 0.66; mean *r* = 0.36; all *P* < 2 × 10^−16^; **Table S2**; **Fig. 2a**). Closer relatives tend to have more similar paternal wealth, and the similarity in paternal wealth between relatives predicts their similarity in occupational status extremely well (Pearson *r* = 0.91; **Fig. 2a; inset**). Thus, there is clear confounding in these data between transmission of genes and the effects of parental wealth on familial similarity in social status. Apart from wealth, numerous other non-genetic factors may contribute to familial correlations (*47, 48*). Two *post hoc* analyses are presented in (29) in an attempt to rule out non-genetic contributors to familial resemblance in social status. In **Supplementary Note 4**, we detail why these analyses are uninformative as to the strength of non-genetic effects on resemblance in social status between relatives (**Fig. S1**).

The confounding of genetic and non-genetic transmission in these data invalidates the interpretation of the model parameters offered in (*30*) as pointing to identifiable genetic contributions (**Supplementary Note 1**). In particular, in the presence of such confounding, the interpretation of *G* and *E* in **Eq. 1** as transmissible genetic (heritable) and random non-genetic effects on a phenotype, respectively, no longer holds. Instead, they can be interpreted as a transmissible component and a random, non-transmissible component. Consequently, the parameter interpreted in (*30*) as narrow-sense heritability, *h*^*2*^, is in fact an estimate of the “total transmissibility” of a trait, *t*^2^, the proportion of trait variance attributable to an unknown compound of transmissible influences on the traits, including genes, culture, wealth, environment, etc. (*10, 12*). The second key parameter, *m*, which (*30*) interpreted as the “spousal correlation in the underlying genetics,” does not represent a genetic correlation between mates. It is instead the spousal correlation in the transmissible component of the trait. *m* is derived from the “intergenerational persistence rate,” 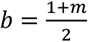, estimated from the regression model. The expected correlation for a given kinship pair is equal to *t*^2^*b*^*n*^, where *n* denotes genealogical distance (**Fig. 1a**). Note that the parameterization of *b* for father-son and grandparent-grandchild relationships also depends on the degree of assortative mating with respect to the focal trait itself; see **Supplementary Note 1**. The conflation of genetic and non-genetic transmission helps to explain why the model parameters estimated in (*30*), which are claimed to represent quantitative genetic parameters *h*^*2*^ and *m*, are much higher than estimates of these same parameters from studies that attempt to account for confounding (e.g. (*19, 49, 50*)).

Conclusions in (*30*) about the insensitivity of social standing to policy and sociopolitical context rest on the similarity of estimates of the parameter *b* across status measures and across time. The claim (*30*) that this stability is due to strong assortative mating on a genetic factor for “social ability” does not hold, given that both genetic and non-genetic factors are transmitted within families. The estimate of *m* tells us nothing about genetic versus non-genetic contributions to assortment, and *b* tells us nothing about the cause of within-family persistence of social status (**Fig. 1a; Supplementary Note 1**).

Regardless of whether due to genetic causes or not, a striking statement from (*30*) is that “*The vast social changes in England since the Industrial Revolution, including mass public schooling, have not increased, in any way, underlying rates of social mobility*”. In point of fact, we found that the estimates of familial correlations in (*30*), and, in turn, estimates of the persistence rate, are heavily affected by statistical artifacts (**Fig. 2b; Supplementary Note 6**). Furthermore, we show that across status measures, parent-offspring correlations — an established measure of social mobility (*51, 52*) — generally decrease over time (**Fig. 2c; Supplementary Note 7**). How could the new measure, “persistence rate”, used by (*30*) lead to such contrasting conclusions? (*30*) offers neither justification for why this measure reflects social mobility, nor explanation for the discrepancies with established measures of mobility used in other literature (e.g., *53*) and applied to the same data.

Some readers have already taken arguments in (*30*) as compelling evidence that social status is largely caused by genetic factors (*54–58*). Yet the assumptions and interpretations in (*30*) ignore a century of quantitative-genetic theory, previous empirical evidence for confounding, and the fallacies that arise when confounding is ignored (*13, 17, 21, 22, 26, 48, 59–66*), as well as patterns in the paper’s own data that conflict with the interpretations presented. In this regard, we emphasize that (*30*) does not merely overstate the findings: the model parameters are misconstrued and the pervasive confounding of genetic and non-genetic transmission is not addressed.

### Are modern genomic studies less susceptible to confounding?

In relying solely on observational phenotypic data and assuming that transmission in families is solely genetic, (*30*) is similar in spirit to studies carried out by Francis Galton a century and a half ago. One might hope that the inferential flaws described above are remedied in studies that use large genomic datasets and employ state-of-the-art statistical methods to adjust for confounding. As we outline, however, the same concerns remain broadly applicable; confounding is still poorly understood and often underplayed in the literature.

#### Confounding in genomic studies is poorly understood

Human geneticists have long appreciated that there are myriad ways by which a genetic variant may be associated with a trait or outcome (*26, 67, 68*). A key example is “population stratification” in genomic data (e.g., in GWAS) wherein patterns of genetic similarity in a sample are correlated with the phenotype studied (**Fig. 1b**). Possible reasons for this correlation include social, environmental (**Box 1-iii**), or genetic (**Box 1-ii**) factors, contemporary and historical. Typically, the specific causes are unknown. These same axes of genetic similarity (“population structure”) are reflected in the frequencies of numerous genetic markers that may be tested for association with a trait in a GWAS. Consequently, any such markers will tend to be correlated with the trait, even if only a subset (or in fact none) of the variants causally affect it (**Fig. 1b**).

Consider, for example, a GWAS conducted to identify genetic risk factors for asthma in a sample of people from the US of either primarily European American genetic ancestry or primarily African American genetic ancestry. There are many millions of variants in the genome that significantly differ in frequency between these groups. At the same time, African Americans in the US are systematically exposed to higher levels of air pollution (*69*), an environmental risk factor for asthma. If confounding is not adequately addressed, the GWAS would lead us to conclude erroneously that “African American genetics” predispose one for asthma (**Box 1-iii**). Regardless of what drives population stratification, it can result in biased estimates of the individual effects of numerous genetic markers that tag these axes of population structure (**Fig. 1b**).

Human geneticists use various methods to adjust for confounded associations. However, confounding may persist, despite application of these methods (“residual confounding”). In 2019, we and other researchers discovered that genetic effect estimates in the largest GWASs for height—the most extensively studied polygenic trait—were biased due to residual confounding (*64, 65*). It is plausible that this confounding is, at least in part, “genetic confounding”: the effect estimated for each individual variant was biased by the effects of many other variants (**Box 1-iii**). Regardless of the source of confounding, it became clear that while the bias for each individual genetic variant was slight, it was systematic across variants. Consequently, when researchers summed over signals from many genetic variants, they also summed over systematic biases. This led to erroneous conclusions in many studies (as detailed in *17, 64, 65*). Further research has suggested that residual confounding may affect many GWASs, in particular for social outcomes and traits that are heavily influenced by social context (*26, 66, 68, 70–72*)

Even so, a lack of statistical identifiability is often not explicitly demonstrable, and it is often difficult to pinpoint the specific confounders in a study. To our knowledge, no general method exists for the detection and quantification of confounding in GWAS data. In the absence of an omnibus litmus test for confounding, the answer cannot be to reduce the burden of proof for causal narratives.

#### Confounding in genomic studies is downplayed

Studies often imply that confounding is completely remedied by current methods, despite evidence to the contrary (*64–66, 68, 70, 72*– *81*). Some methods to estimate genetic parameters have grown in popularity even after they were shown to be susceptible to confounding, with this susceptibility rarely mentioned as a caveat (see, e.g., discussions in *64, 71, 72, 76, 82, 83*). In other cases, confounding is acknowledged as a potential limitation, but its impact on the reported results (and their interpretation) is downplayed or obscured (see, for instance, (*84, 85*)).

As one example, consider the reporting of evidence for genetic effects from standard GWASs versus family-based studies. Family studies identify genotype-trait associations within, instead of among, families. This approach greatly mitigates many sources of confounding (*66, 68, 86*). Family studies have yielded estimates of genetic effects on behavior or social outcomes that are substantially weaker than those estimated from standard GWASs (*49, 66, 70, 72, 87–90*). Reporting practices tend to downplay this point by using evidence from family studies as support for *existence* of a causal genetic effect, while continuing to rely on the magnitude of effects estimated in a standard GWAS (*61*). Such reporting choices mislead by presenting signals susceptible to confounding as measures of genetic causality.

#### Confounding in complex traits genetics: aggregation of many small biases

Quantitative geneticists acknowledge residual confounding as an unsolved problem. But in practice, researchers face incentives to publish their inferences of genetic associations that are vulnerable to confounding. With polygenic (or “complex”) traits, the usual focus of these studies, genetic contributions to trait variation are largely due to numerous genetic variants with individually small effects. Researchers often wish to leverage weaker and weaker genetic associations to capture these highly polygenic signals. At the same time, confounding tends to be aggravated as more weakly associated variants are considered (*66, 80*). Thus, in the pursuit of understanding polygenic effects, researchers may face a tradeoff between explaining a smaller part of the phenomenon under study in a causally rigorous way, versus accounting for a seemingly larger part, at the price of unknown biases introduced by confounding.

An example of this tradeoff lies in genetic trait prediction with so-called “polygenic scores” (*91*). Polygenic scores based on more variants, including weakly-associated ones, may be preferred by researchers because they often attain higher prediction accuracy than polygenic scores that are limited to confident associations. However, polygenic scores that include many weakly-associated variants are plausibly more susceptible to underappreciated axes of confounding (*66, 80, 92*). Subsequent “consumers”, including clinicians, researchers, policymakers, and the general public, may then assume these polygenic scores capture strictly direct genetic effects; the possibility of confounding is rarely acknowledged. Consider, for instance, an hypothetical preimplantation genetic testing using a polygenic score based on the asthma GWAS we described above. In this extreme, embryos would mistakenly be prioritized for implantation according to whether or not they share genetic variants with people exposed to higher levels of air pollution in a previous generation.

Similarly, popular methods to estimate genetic correlations (the correlation between two groups of individuals in genetic effects on a trait, or the correlation of genetic effects on two traits) often indiscriminately aggregate across genome-wide associations (*93, 94*). Such methods are useful for characterizing how the genetic bases of complex traits are intertwined. However, they may inadvertently mask (or even introduce) additional axes of confounding. For example, confounding by population stratification that is shared between the two groups or two traits, or confounding with genetic effects on a third trait, can lead to biases in genetic correlation estimates (*66, 76, 82, 95, 96*). The use of genetic correlations in causal inference (e.g., in “Mendelian Randomization” (*97–99*)) therefore remains controversial (*76, 100*). Yet when a study reports conclusions based on *genetic correlations*, it is likely to be interpreted—particularly by non-experts—as unambiguously reflecting genetic causality.

## Further confounding via research choices

Such unknown axes of confounding are plausibly a concern in a recent study that, based on an analysis of genetic correlations, purported to resolve an evolutionary paradox: why alleles associated with same-sex sexual behavior are maintained, despite being “reproductively disadvantageous” (*31*). In what follows, we focus on what we refer to as “analytical confounding,” where potential confounding is worsened, or even introduced, through researchers’ analysis choices. In (*31*): confusing model assumptions with evidence, ignoring the compatibility of data with confounded explanations, and introducing confounding through researchers’ classification choices. We posit that, while these problems are not unique to genomic studies, they can evade attention when couched in reports about how behaviors and outcomes are *genetically correlated*.

### Confusion of assumptions with evidence

In (*31*), a measure of bisexual behavior is defined based on questionnaire data about total lifetime number of sexual partners and same-sex sexual partners (hereafter, we refer to this measure as BSB; see (*101*) and (*102*) for discussion of shortcomings of such measures). The paper reports a significant positive genetic correlation between BSB in males and the number of children. But when adjusting this genetic correlation for genetic correlations of each measure with self-assessment as a “risk-taker”, the adjusted (or “partial”) genetic correlation between BSB in males and number of children was statistically indistinguishable from zero (**Fig. 3a**). It presents an interpretation of this finding as evidence that “the current genetic maintenance of male BSB is a by-product of selection for male risk-taking behavior.” No explanation of the hypothesized mechanism by which risk-taking behavior increases the number of offspring is given in (*31*), but in subsequent news coverage, one of the authors is quoted, “self-reported risk-taking [likely] includes unprotected sex and promiscuity, which could result in more children” (*103*).

**Figure 3.**
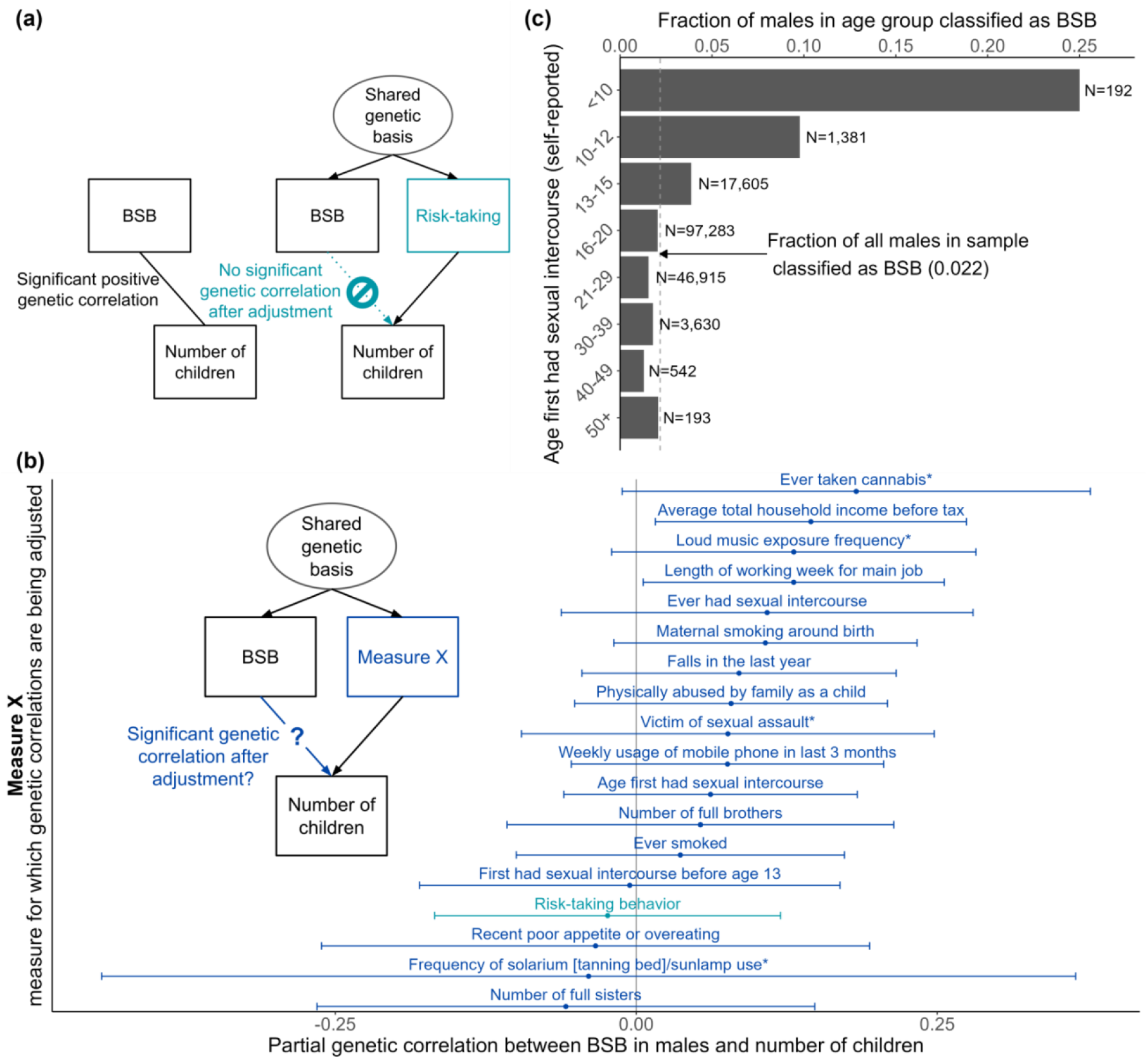
**(a)** Song and Zhang (2024) show that the estimated genetic correlation between BSB (a measure of bisexual behavior) in males and number of children is significantly different from zero (left diagram). They hypothesized the causal structure shown in the right diagram: Genetic variants affecting BSB affect the number of children only through their simultaneous effect on risk-taking behavior. When adjusting for genetic effects on risk-taking behavior, the residualized (or “partial”) genetic correlation between BSB and number of children is no longer significantly nonzero. They take this observation as evidence for their hypothesis. **(b)** When we repeat this analysis but replace risk taking with a variety of other measures (blue in causal diagram), 16/18 measures yield a partial genetic correlation between BSB in males and number of children that is consistent with zero (measures we considered are shown in blue in the plot on the right; error bars indicate 95% confidence intervals). Asterisks indicate the four measures that have no significant partial genetic correlation with number of children (Fig S16). **(c)** Male participants in the study sample who reported having first had sex before age 13 (including victims of childhood sexual assault, many of which would have had same-sex perpetrators) are likelier to be classified as BSB by the criteria used in Song and Zhang (2024). N, total number of males in each group.

The study presents the contrast between these unadjusted and partial genetic correlations as support for a causal claim. However, the causal model is assumed *a priori*, and no evidence supporting this model is provided. Even under the assumption that the three measures considered are the only ones at play—and some causally affect others—the evidence is equally consistent with contradictory causal hypotheses (e.g., different directions of causality, arrows in Fig. 2b of and **Figs. 3a, S14** here; **Supplementary Note 8**; c.f. (*104*)).

Furthermore, the study did not evaluate the support for any alternative model involving other factors, observed or latent. The authors justify their focus on risk-taking as a mechanistic explanation for the genetic maintenance of same-sex sexual behavior by citing previous reports of genetic correlations of same-sex sexual behavior and risk-taking (*105, 106*). However, these studies (and others (*107*)) reported multiple measures with similar (or even stronger) genetic correlations with same-sex sexual behavior than risk-taking (*105*) (**Supplementary Table 5**). Sexual behavior aside, (*31*) neither cites nor offers any evidence for the alleles associated with risk-taking being maintained over a long evolutionary timescale. Additionally, this association is based on an answer to a single questionnaire question, “Would you describe yourself as someone who takes risks?” (*108*). It is possible that responses to this question reflect a tendency towards practicing unprotected sex and promiscuity, and that they simultaneously correlate with risk-taking tendencies that have been relevant for fitness throughout recent human evolution and across evolving societies; but, as acknowledged in (*31*), these key assumptions are hard to evaluate.

For these reasons, we asked: Is there unique support for the assumed mechanistic model, in particular the role of risk-taking behavior as a mediator? To answer this question, we considered models wherein a measure other than risk-taking mediates the genetic correlation between BSB and number of children (“Measure X” in blue in **Fig. 3b**). If adjusting for this measure also results in a partial genetic correlation between BSB and number of children that is not significantly different from 0, the data are equally compatible with the hypothesis that there is a reproductive advantage for BSB-affecting alleles because of their simultaneous effect on Measure X. We implemented this strategy with 18 measures, selected based on prior evidence of high genetic correlations with same-sex sexual behavior, risk-taking behavior, and/or number of children (*107*) (**Supplementary Tables 4-5**; **Supplementary Note 8**). All but two of these models yielded a partial genetic correlation between BSB and number of children that was not significantly different from 0 (Genomic SEM (*109*) P >0.05 before applying any correction for multiple testing) (**Fig. 3b**). Hence, other causal narratives that do not involve risk taking could just as easily be constructed: the data are equally consistent with the hypothesis that genetic variants driving BSB are maintained through evolution as a byproduct of selection on the number of falls in the past year, usage of cell phone, or any of these measures (**Fig. 3b; Fig. S15**). The ease with which alternative hypotheses about genetic causality receive support using the method from (*31*) is revealing: the conclusion is highly reliant on the choice of a single hypothesis to test and a failure to consider confounding (**Box 1-iv**).

### Confounding introduced by researchers’ classification choices

Measures relating to having experienced some form of nonconsensual sex (“victim of sexual assault” and “first had sex before age 13”) exhibited some of the strongest genetic correlations with BSB in males (**Fig. S15**). This observation led us to be concerned about the ascertainment choices made in (*31*). Indeed, we found that classification as a BSB individual is highly enriched among males who reported having first had sex before age 13 (in this regard, we note that children under this age are not legally capable of consenting to any sexual activity in the UK (*110*)) (**Fig. 3c**). Whereas 2.2% of males in the sample considered were classified as “BSB individuals,” this classification rate increased to 9.8% among those who reported first having sex between ages 10-12 (inclusively), and 25% among those who reported first having had sex before age 10. For this dataset, there is no information about the age at which males classified as BSB first had same-sex sexual intercourse, or what fraction of victims of sexual assault had a perpetrator of the same sex. It is, however, established that the majority of reported sexual assaults on prepubescent male victims are carried out by male perpetrators (*111–114*). This aggravates the concern that the BSB classification used in (*31*) conflated voluntary sexual behavior and sexual assault, undermining the study’s stated aim of advancing our understanding of human sexual preferences. Taken together, our reanalysis of (*31*) cautions yet again against causal inference based on preferential attention towards sensational hypotheses and analyses that seemingly support them.

### Conclusion

The study of the genetics underlying human behavior and social outcomes, with its fraught history and heightened potential for misinterpretation and misappropriation (*61, 62, 71, 101, 115, 116*), demands the utmost rigor. The failure to reckon with confounding fuels misinterpretation of genetics research and impedes scientific progress. We are therefore concerned that a publishing culture that rewards sensationalism may instead promote a decline in standards (*117, 118*). In that respect, everyone has a role to play: it is crucial that researchers, reviewers, and editors uphold the highest standards in their handling of these complex, far-reaching issues.

## Acknowledgements

We thank Ipsita Agarwal, Kate Antonovics, Mark Borrello, Raj Chetty, Graham Coop, Doc Edge, Sasha Gusev, Kelley Harris, Mark Kirkpatrick, Magnus Nordborg, Nick Patterson, Molly Przeworski, Sohini Ramachandran, Noah Rosenberg, James Schmitz, Elizabeth Thompson, Elliot Tucker-Drob, Alex Viloria-Winnett and the Biological Interest Group of the Minnesota Center for Philosophy of Science for comments on the manuscript and helpful discussions. We thank Gregory Clark for providing us with a corrected version of the occupational status data. We thank Siliang Song and Jianzhi (George) Zhang for providing us with detailed methods to replicate the GWAS they had performed. This research has been conducted using the UK Biobank Resource under Application Number 92741. The work was funded by NSF grant DBI-2010892 to J.W.B, NSF Graduate Research Fellowship DGE 2137420 to O.S.S., and NIH grant R35GM151108 and a Pew Scholarship to A.H.

## Data availability

All code for reproducing our analyses is available at https://github.com/harpak-lab/confounding.

## References

1. S. F. Galton, Hereditary Genius: An Inquiry Into Its Laws and Consequences (D. Appleton, 1869).

2. B. Mehler, Hereditarianism, The Wiley Blackwell Encyclopedia of Race, Ethnicity, and Nationalism (2015) pp. 1–3.

3. S. Wright, Statistical Methods in Biology. J. Am. Stat. Assoc. 26, 155–163 (1931).

4. L. L. Cavalli-Sforza, M. W. Feldman, Models for cultural inheritance I. Group mean and within group variation. Theor. Popul. Biol. 4, 42–55 (1973).

5. L. L. Cavalli-Sforza, M. W. Feldman, Cultural versus biological inheritance: phenotypic transmission from parents to children. (A theory of the effect of parental phenotypes on children’s phenotypes). Am. J. Hum. Genet. 25, 618–637 (1973).

6. D. C. Rao, N. E. Morton, S. Yee, Analysis of family resemblance. II. A linear model for familial correlation. Am. J. Hum. Genet. 26, 331–359 (1974).

7. D. C. Rao, N. E. Morton, S. Yee, Resolution of cultural and biological inheritance by path analysis. Am. J. Hum. Genet. 28, 228–242 (1976).

8. L. L. Cavalli-Sforza, M. W. Feldman, The Evolution of Continuous Variation. III. Joint Transmission of Genotype, Phenotype and Environment. Genetics 90, 391–425 (1978).

9. J. Rice, C. R. Cloninger, T. Reich, Multifactorial inheritance with cultural transmission and assortative mating. I. Description and basic properties of the unitary models. Am. J. Hum. Genet. 30, 618–643 (1978).

10. C. R. Cloninger, J. Rice, T. Reich, Multifactorial inheritance with cultural transmission and assortative mating. II. a general model of combined polygenic and cultural inheritance. Am. J. Hum. Genet. 31, 176–198 (1979).

11. C. R. Cloninger, J. Rice, T. Reich, Multifactorial inheritance with cultural transmission and assortative mating. III. Family structure and the analysis of separation experiments. Am. J. Hum. Genet. 31, 366–388 (1979).

12. J. Rice, C. R. Cloninger, T. Reich, Analysis of behavioral traits in the presence of cultural transmission and assortative mating: Applications to IQ and SES. Behav. Genet. 10, 73–92 (1980).

13. R. C. Lewontin, S. P. R. Rose, L. J. Kamin, Not in Our Genes: Biology, Ideology, and Human Nature (Pantheon Books, New York, 1984)vol. 152.

14. B. J. Vilhjálmsson, M. Nordborg, The nature of confounding in genome-wide association studies. Nat. Rev. Genet. 14, 1–2 (2013).

15. M. W. Feldman, F. B. Christiansen, S. P. Otto, Gene-culture co-evolution: teaching, learning, and correlations between relatives. Isr. J. Ecol. Evol. 59, 72–91 (2013).

16. G. Solon, Theoretical models of inequality transmission across multiple generations. Res. Soc. Stratif. Mobil. 35, 13–18 (2014).

17. N. Barton, J. Hermisson, M. Nordborg, Why structure matters. Elife 8 (2019).

18. R. Uchiyama, R. Spicer, M. Muthukrishna, Cultural evolution of genetic heritability. Behav. Brain Sci. 45, e152 (2022).

19. M. D. Collado, I. Ortuño-Ortín, J. Stuhler, Estimating Intergenerational and Assortative Processes in Extended Family Data. Rev. Econ. Stud. 90, 1195–1227 (2023).

20. A. F. Herzig, C. Noûs, A. S. Pierre, H. Perdry, A model for co-occurrent assortative mating and vertical cultural transmission and its impact on measures of genetic associations, bioRxiv (2023)p. 2023.04.08.536101.

21. R. C. Lewontin, Annotation: the analysis of variance and the analysis of causes. Am. J. Hum. Genet. 26, 400–411 (1974).

22. M. W. Feldman, R. C. Lewontin, The heritability hang-up. Science 190, 1163–1168 (1975).

23. R. C. Bailey, Hereditarian scientific fallacies. Genetica 99, 125–133 (1997).

24. N. A. Holtzman, Genetics and social class. J. Epidemiol. Community Health 56, 529–535 (2002).

25. M. W. Feldman, S. Ramachandran, Missing compared to what? Revisiting heritability, genes and culture. Philos. Trans. R. Soc. Lond. B Biol. Sci. 373 (2018).

26. A. I. Young, S. Benonisdottir, M. Przeworski, A. Kong, Deconstructing the sources of genotype-phenotype associations in humans. Science 365, 1396–1400 (2019).

27. H. Shen, M. W. Feldman, Drowning in shallow causality. Behav. Brain Sci. 46, e199 (2023).

28. A. S. Goldberger, Heritability. Economica 46, 327–347 (1979).

29. K. N. Lala, M. W. Feldman, Genes, culture, and scientific racism. Proc. Natl. Acad. Sci. U. S. A. 121, e2322874121 (2024).

30. G. Clark, The inheritance of social status: England, 1600 to 2022. Proc. Natl. Acad. Sci. U. S. A. 120, e2300926120 (2023).

31. S. Song, J. Zhang, Genetic variants underlying human bisexual behavior are reproductively advantageous. Sci Adv 10, eadj6958 (2024).

32. S. Greenland, J. Pearl, J. M. Robins, Causal diagrams for epidemiologic research. Epidemiology 10, 37–48 (1999).

33. M. A. Hernan, J. M. Robins, Causal Inference: What If (CRC Press, Boca Raton, FL, 2025).

34. J. Pearl, Causality: Models, Reasoning, and Inference (Cambridge University Press, Cambridge, England, 2009).

35. T. J. Vanderweele, Surrogate measures and consistent surrogates. Biometrics 69, 561–569 (2013).

36. J. Pearl, Causal diagrams for empirical research. Biometrika 82, 669 (1995).

37. R. A. Fisher, The Correlation between Relatives on the Supposition of Mendelian Inheritance. Trans. R. Soc. Edinb. 52, 399–433 (1918).

38. A. Gimelfarb, A general linear model for the genotypic covariance between relatives under assortative mating. J. Math. Biol. 13, 209–226 (1981).

39. G. Clark, The Son Also Rises (Princeton University Press, 2014).

40. G. Clark, As a hereditarian, I strongly support economic redistribution, Quillette (2023). https://quillette.com/2023/08/21/hereditarianism-and-economic-redistribution/.

41. F. J. Odling-Smee, “Niche-constructing phenotypes” in The Role of Behavior in Evolution, (pp, H. C. Plotkin, Ed. (The MIT Press, viii, Cambridge, MA, US, 1988)vol. 198, pp. 73–132.

42. M. Hällsten, F. T. Pfeffer, Grand Advantage: Family Wealth and Grandchildren’s Educational Achievement in Sweden. Am. Sociol. Rev. 82, 328–360 (2017).

43. Rising Inequality, Schools, and Children’s Life Chances (Russell Sage Foundation, 2011).

44. E. Karagiannaki, The effect of parental wealth on children’s outcomes in early adulthood. J. Econ. Inequality 15, 217–243 (2017).

45. R. K. Q. Akee, W. E. Copeland, G. Keeler, A. Angold, E. J. Costello, Parents’ Incomes and Children’s Outcomes: A Quasi-experiment Using Transfer Payments from Casino Profits. Am. Econ. J. Appl. Econ. 2, 86–115 (2010).

46. K. Cooper, K. Stewart, Does Household Income Affect children’s Outcomes? A Systematic Review of the Evidence. Child Indic. Res. 14, 981–1005 (2021).

47. M. Feldman, L. L. Cavalli-Sforza, Cultural Transmission and Evolution: A Quantitative Approach (Princeton University Press, 1981).

48. E. Turkheimer, Three Laws of Behavior Genetics and What They Mean. Curr. Dir. Psychol. Sci. 9, 160–164 (2000).

49. A. I. Young, M. L. Frigge, D. F. Gudbjartsson, G. Thorleifsson, G. Bjornsdottir, P. Sulem, G. Masson, U. Thorsteinsdottir, K. Stefansson, A. Kong, Relatedness disequilibrium regression estimates heritability without environmental bias. Nat. Genet. 50, 1304–1310 (2018).

50. L. Yengo, M. R. Robinson, M. C. Keller, K. E. Kemper, Y. Yang, M. Trzaskowski, J. Gratten, P. Turley, D. Cesarini, D. J. Benjamin, N. R. Wray, M. E. Goddard, J. Yang, P. M. Visscher, Imprint of assortative mating on the human genome. Nat Hum Behav 2, 948–954 (2018).

51. O. Causa, Å. Johansson, “Intergenerational Social Mobility” (Organisation for Economic Co-Operation and Development (OECD), 2009); 10.1787/223106258208.

52. R. Chetty, N. Hendren, P. Kline, E. Saez, Where is the land of Opportunity? The Geography of Intergenerational Mobility in the United States. Q. J. Econ. 129, 1553–1623 (2014).

53. A. Miles, Social Mobility in Nineteenth- and Early Twentieth-Century England (Palgrave Macmillan UK, 1999).

54. J. J. Lee, The heritability and persistence of social class in England. Proc. Natl. Acad. Sci. U. S. A. 120, e2309250120 (2023).

55. C. Cosh, Colby Cosh: Is socioeconomic status hereditary?, National Post (2023). https://nationalpost.com/opinion/is-socioeconomic-status-hereditary.

56. G. N. Marks, Has Cognitive Ability Become More Important for Education and the Labor Market? A Comparison of the Project Talent and 1979 National Longitudinal Survey of Youth Cohorts. J Intell 11 (2023).

57. J. Ratia, El Estatus Social También se Hereda, Ethic (2024). https://ethic.es/2024/01/el-estatus-social-tambien-se-hereda/.

58. O. Mayo, V. Nanjundiah, Reflections on assortative mating, social stratification, and genetics. J. Genet. 103, 15 (2024).

59. S. Scarr, K. McCartney, How people make their own environments: A theory of genotype -- > environment effects. Child Dev. 54, 424 (1983).

60. M. Feldman, Echoes of the past: hereditarianism and A Troublesome Inheritance. PLoS Genet. 10, e1004817 (2014).

61. G. Coop, M. Przeworski, Lottery, luck, or legacy. A review of “The Genetic Lottery: Why DNA matters for social equality.” Evolution 76, 846–853 (2022).

62. G. Coop, M. Przeworski, Luck, lottery, or legacy? The problem of confounding. A reply to Harden. Evolution 76, 2464–2468 (2022).

63. A. S. Young, Estimation of indirect genetic effects and heritability under assortative mating, bioRxiv (2023)p. 2023.07.10.548458.

64. J. J. Berg, A. Harpak, N. Sinnott-Armstrong, A. M. Joergensen, H. Mostafavi, Y. Field, E. A. Boyle, X. Zhang, F. Racimo, J. K. Pritchard, G. Coop, Reduced signal for polygenic adaptation of height in UK Biobank. Elife 8 (2019).

65. M. Sohail, R. M. Maier, A. Ganna, A. Bloemendal, A. R. Martin, M. C. Turchin, C. W. K. Chiang, J. Hirschhorn, M. J. Daly, N. Patterson, B. Neale, I. Mathieson, D. Reich, S. R. Sunyaev, Polygenic adaptation on height is overestimated due to uncorrected stratification in genome-wide association studies. Elife 8, e39702 (2019).

66. H. Mostafavi, A. Harpak, I. Agarwal, D. Conley, J. K. Pritchard, M. Przeworski, Variable prediction accuracy of polygenic scores within an ancestry group. Elife 9 (2020).

67. E. S. Lander, N. J. Schork, Genetic dissection of complex traits. Science 265, 2037–2048 (1994).

68. C. Veller, G. M. Coop, Interpreting population- and family-based genome-wide association studies in the presence of confounding. PLoS Biol. 22, e3002511 (2024).

69. K. K. Nishimura, J. M. Galanter, L. A. Roth, S. S. Oh, N. Thakur, E. A. Nguyen, S. Thyne, H. J. Farber, D. Serebrisky, R. Kumar, E. Brigino-Buenaventura, A. Davis, M. A. LeNoir, K. Meade, W. Rodriguez-Cintron, P. C. Avila, L. N. Borrell, K. Bibbins-Domingo, J. R. Rodriguez-Santana, Ś. Sen, F. Lurmann, J. R. Balmes, E. G. Burchard, Early-life air pollution and asthma risk in minority children. The GALA II and SAGE II studies. Am. J. Respir. Crit. Care Med. 188, 309–318 (2013).

70. A. Okbay, Y. Wu, N. Wang, H. Jayashankar, M. Bennett, S. M. Nehzati, J. Sidorenko, H. Kweon, G. Goldman, T. Gjorgjieva, Y. Jiang, B. Hicks, C. Tian, D. A. Hinds, R. Ahlskog, P. K. E. Magnusson, S. Oskarsson, C. Hayward, A. Campbell, D. J. Porteous, J. Freese, P. Herd, 23andMe Research Team, Social Science Genetic Association Consortium, C. Watson, J. Jala, D. Conley, P. D. Koellinger, M. Johannesson, D. Laibson, M. N. Meyer, J. J. Lee, A. Kong, L. Yengo, D. Cesarini, P. Turley, P. M. Visscher, J. P. Beauchamp, D. J. Benjamin, A. I. Young, Polygenic prediction of educational attainment within and between families from genome-wide association analyses in 3 million individuals. Nat. Genet. 54, 437–449 (2022).

71. M. N. Meyer, P. S. Appelbaum, D. J. Benjamin, S. L. Callier, N. Comfort, D. Conley, J. Freese, N. A. Garrison, E. M. Hammonds, K. P. Harden, S. S.-J. Lee, A. R. Martin, D. O. Martschenko, B. M. Neale, R. H. C. Palmer, J. Tabery, E. Turkheimer, P. Turley, E. Parens, Wrestling with Social and Behavioral Genomics: Risks, Potential Benefits, and Ethical Responsibility. Hastings Cent. Rep. 53 Suppl 1, S2–S49 (2023).

72. M. G. Nivard, D. W. Belsky, K. P. Harden, T. Baier, O. A. Andreassen, E. Ystrøm, E. van Bergen, T. H. Lyngstad, More than nature and nurture, indirect genetic effects on children’s academic achievement are consequences of dynastic social processes. Nat Hum Behav, doi: 10.1038/s41562-023-01796-2 (2024).

73. G. Sella, N. H. Barton, Thinking About the Evolution of Complex Traits in the Era of Genome-Wide Association Studies. Annu. Rev. Genomics Hum. Genet. 20, 461–493 (2019).

74. A. A. Zaidi, I. Mathieson, Demographic history mediates the effect of stratification on polygenic scores. Elife 9 (2020).

75. H. Wang, B. Aragam, E. Xing, Tradeoffs of Linear Mixed Models in Genome-wide Association Studies, arXiv [q-bio.QM] (2021). http://arxiv.org/abs/2111.03739.

76. R. Border, G. Athanasiadis, A. Buil, A. J. Schork, N. Cai, A. I. Young, T. Werge, J. Flint, K. S. Kendler, S. Sankararaman, A. W. Dahl, N. A. Zaitlen, Cross-trait assortative mating is widespread and inflates genetic correlation estimates. Science 378, 754–761 (2022).

77. A. I. Young, S. M. Nehzati, S. Benonisdottir, A. Okbay, H. Jayashankar, C. Lee, D. Cesarini, D. J. Benjamin, P. Turley, A. Kong, Mendelian imputation of parental genotypes improves estimates of direct genetic effects. Nat. Genet. 54, 897–905 (2022).

78. M. Onifade, M.-H. Roy-Gagnon, M.-É. Parent, K. M. Burkett, Comparison of mixed model based approaches for correcting for population substructure with application to extreme phenotype sampling. BMC Genomics 23, 98 (2022).

79. Y. Yao, A. Ochoa, Limitations of principal components in quantitative genetic association models for human studies. Elife 12 (2023).

80. A. J. Aw, J. McRae, E. Rahmani, Y. S. Song, Highly parameterized polygenic scores tend to overfit to population stratification via random effects, bioRxiv (2024)p. 2024.01.27.577589.

81. K. E. Grinde, B. L. Browning, A. P. Reiner, T. A. Thornton, S. R. Browning, Adjusting for principal components can induce spurious associations in genome-wide association studies in admixed populations, bioRxiv (2024)p. 2024.04.02.587682.

82. S. Zabad, A. P. Ragsdale, R. Sun, Y. Li, S. Gravel, Assumptions about frequency-dependent architectures of complex traits bias measures of functional enrichment. Genet. Epidemiol. 45, 621–632 (2021).

83. N. LaPierre, B. Fu, S. Turnbull, E. Eskin, S. Sankararaman, Leveraging family data to design Mendelian randomization that is provably robust to population stratification. Genome Res. 33, 1032–1041 (2023).

84. E. J. Giangrande, E. Turkheimer, Race, Ethnicity, and the Scarr-Rowe Hypothesis: A Cautionary Example of Fringe Science Entering the Mainstream. Perspect. Psychol. Sci. 17, 696–710 (2022).

85. E. Turkheimer, S. R. Greer, Spit for Science and the Limits of Applied Psychiatric Genetics. Philos. Psychiatr. Psychol., doi: 10.1353/ppp.0.a923702 (2024).

86. R. S. Spielman, R. E. McGinnis, W. J. Ewens, Transmission test for linkage disequilibrium: the insulin gene region and insulin-dependent diabetes mellitus (IDDM). Am. J. Hum. Genet. 52, 506–516 (1993).

87. J. J. Lee, R. Wedow, A. Okbay, E. Kong, O. Maghzian, M. Zacher, T. A. Nguyen-Viet, P. Bowers, J. Sidorenko, R. Karlsson Linnér, M. A. Fontana, T. Kundu, C. Lee, H. Li, R. Li, R. Royer, P. N. Timshel, R. K. Walters, E. A. Willoughby, L. Yengo, 23andMe Research Team, COGENT (Cognitive Genomics Consortium), Social Science Genetic Association Consortium, M. Alver, Y. Bao, D. W. Clark, F. R. Day, N. A. Furlotte, P. K. Joshi, K. E. Kemper, A. Kleinman, C. Langenberg, R. Mägi, J. W. Trampush, S. S. Verma, Y. Wu, M. Lam, J. H. Zhao, Z. Zheng, J. D. Boardman, H. Campbell, J. Freese, K. M. Harris, C. Hayward, P. Herd, M. Kumari, T. Lencz, J. ‘an Luan, A. K. Malhotra, A. Metspalu, L. Milani, K. K. Ong, J. R. B. Perry, D. J. Porteous, M. D. Ritchie, M. C. Smart, B. H. Smith, J. Y. Tung, N. J. Wareham, J. F. Wilson, J. P. Beauchamp, D. C. Conley, T. Esko, S. F. Lehrer, P. K. E. Magnusson, S. Oskarsson, T. H. Pers, M. R. Robinson, K. Thom, C. Watson, C. F. Chabris, M. N. Meyer, D. I. Laibson, J. Yang, M. Johannesson, P. D. Koellinger, P. Turley, P. M. Visscher, D. J. Benjamin, D. Cesarini, Gene discovery and polygenic prediction from a genome-wide association study of educational attainment in 1.1 million individuals. Nat. Genet. 50, 1112–1121 (2018).

88. S. Trejo, B. W. Domingue, Genetic nature or genetic nurture? Introducing social genetic parameters to quantify bias in polygenic score analyses. Biodemography Soc. Biol. 64, 187–215 (2018).

89. S. Selzam, S. J. Ritchie, J.-B. Pingault, C. A. Reynolds, P. F. O’Reilly, R. Plomin, Comparing Within- and Between-Family Polygenic Score Prediction. Am. J. Hum. Genet. 105, 351–363 (2019).

90. L. J. Howe, M. G. Nivard, T. T. Morris, A. F. Hansen, H. Rasheed, Y. Cho, G. Chittoor, R. Ahlskog, P. A. Lind, T. Palviainen, M. D. van der Zee, R. Cheesman, M. Mangino, Y. Wang, S. Li, L. Klaric, S. M. Ratliff, L. F. Bielak, M. Nygaard, A. Giannelis, E. A. Willoughby, C. A. Reynolds, J. V. Balbona, O. A. Andreassen, H. Ask, A. Baras, C. R. Bauer, D. I. Boomsma, A. Campbell, H. Campbell, Z. Chen, P. Christofidou, E. Corfield, C. C. Dahm, D. R. Dokuru, L. M. Evans, E. J. C. de Geus, S. Giddaluru, S. D. Gordon, K. P. Harden, W. D. Hill, A. Hughes, S. M. Kerr, Y. Kim, H. Kweon, A. Latvala, D. A. Lawlor, L. Li, K. Lin, P. Magnus, P. K. E. Magnusson, T. T. Mallard, P. Martikainen, M. C. Mills, P. R. Njølstad, J. D. Overton, N. L. Pedersen, D. J. Porteous, J. Reid, K. Silventoinen, M. C. Southey, C. Stoltenberg, E. M. Tucker-Drob, M. J. Wright, J. K. Hewitt, M. C. Keller, M. C. Stallings, J. J. Lee, K. Christensen, S. L. R. Kardia, P. A. Peyser, J. A. Smith, J. F. Wilson, J. L. Hopper, S. Hägg, T. D. Spector, J.-B. Pingault, R. Plomin, A. Havdahl, M. Bartels, N. G. Martin, S. Oskarsson, A. E. Justice, I. Y. Millwood, K. Hveem, Ø. Naess, C. J. Willer, B. O. Åsvold, P. D. Koellinger, J. Kaprio, S. E. Medland, R. G. Walters, D. J. Benjamin, P. Turley, D. M. Evans, G. Davey Smith, C. Hayward, B. Brumpton, G. Hemani, N. M. Davies, Within-sibship genome-wide association analyses decrease bias in estimates of direct genetic effects. Nat. Genet. 54, 581–592 (2022).

91. A. Torkamani, N. E. Wineinger, E. J. Topol, The personal and clinical utility of polygenic risk scores. Nat. Rev. Genet. 19, 581–590 (2018).

92. B. J. Vilhjálmsson, J. Yang, H. K. Finucane, A. Gusev, S. Lindström, S. Ripke, G. Genovese, P.-R. Loh, G. Bhatia, R. Do, T. Hayeck, H.-H. Won, Schizophrenia Working Group of the Psychiatric Genomics Consortium, Discovery, Biology, and Risk of Inherited Variants in Breast Cancer (DRIVE) study, S. Kathiresan, M. Pato, C. Pato, R. Tamimi, E. Stahl, N. Zaitlen, B. Pasaniuc, G. Belbin, E. E. Kenny, M. H. Schierup, P. De Jager, N. A. Patsopoulos, S. McCarroll, M. Daly, S. Purcell, D. Chasman, B. Neale, M. Goddard, P. M. Visscher, P. Kraft, N. Patterson, A. L. Price, Modeling Linkage Disequilibrium Increases Accuracy of Polygenic Risk Scores. Am. J. Hum. Genet. 97, 576–592 (2015).

93. J. Yang, B. Benyamin, B. P. McEvoy, S. Gordon, A. K. Henders, D. R. Nyholt, P. A. Madden, A. C. Heath, N. G. Martin, G. W. Montgomery, M. E. Goddard, P. M. Visscher, Common SNPs explain a large proportion of the heritability for human height. Nat. Genet. 42, 565–569 (2010).

94. B. Bulik-Sullivan, H. K. Finucane, V. Anttila, A. Gusev, F. R. Day, P.-R. Loh, ReproGen Consortium, Psychiatric Genomics Consortium, Genetic Consortium for Anorexia Nervosa of the Wellcome Trust Case Control Consortium 3, L. Duncan, J. R. B. Perry, N. Patterson, E. B. Robinson, M. J. Daly, A. L. Price, B. M. Neale, An atlas of genetic correlations across human diseases and traits. Nat. Genet. 47, 1236–1241 (2015).

95. W. van Rheenen, W. J. Peyrot, A. J. Schork, S. H. Lee, N. R. Wray, Genetic correlations of polygenic disease traits: from theory to practice. Nat. Rev. Genet. 20, 567–581 (2019).

96. T. T. Morris, N. M. Davies, G. Hemani, G. D. Smith, Population phenomena inflate genetic associations of complex social traits. Sci. Adv. 6, eaay0328 (2020).

97. G. Davey Smith, G. Hemani, Mendelian randomization: genetic anchors for causal inference in epidemiological studies. Hum. Mol. Genet. 23, R89–98 (2014).

98. E. Sanderson, M. M. Glymour, M. V. Holmes, H. Kang, J. Morrison, M. R. Munafò, T. Palmer, C. M. Schooling, C. Wallace, Q. Zhao, G. D. Smith, Mendelian randomization. Nat. Rev. Methods Primers 2, 1–21 (2022).

99. R. C. Richmond, G. Davey Smith, Mendelian randomization: Concepts and scope. Cold Spring Harb. Perspect. Med. 12, a040501 (2022).

100. X. Hu, M. Cai, J. Xiao, X. Wan, Z. Wang, H. Zhao, C. Yang, Benchmarking Mendelian randomization methods for causal inference using genome-wide association study summary statistics. Am. J. Hum. Genet. 0 (2024).

101. C. Ventresca, D. O. Martschenko, R. Wedow, M. Civilek, J. Tabery, J. Carlson, S. C. J. Parker, P. S. Ramos, The Methodological and Ethical Concerns of Genetic Studies of Same-Sex Sexual Behavior. Am. J. Hum. Genet. ([in press]. 2024).

102. I. G. Vázquez, The gay gene(s)? Rethinking the concept of sexual orientation in the context of science. Biol. Philos. 37, 45 (2022).

103. K. Tan, M. D. D. Tan, Genetic variants underlying male bisexual behavior, risk-taking linked to more children – study (2024). https://outragemag.com/genetic-variants-underlying-male-bisexual-behavior-risk-taking-linked-to-more-children-study/.

104. J. G. Bullock, D. P. Green, S. E. Ha, Yes, but what’s the mechanism? (don’t expect an easy answer). J. Pers. Soc. Psychol. 98, 550–558 (2010).

105. A. Ganna, K. J. H. Verweij, M. G. Nivard, R. Maier, R. Wedow, A. S. Busch, A. Abdellaoui, S. Guo, J. F. Sathirapongsasuti, 23andMe Research Team, P. Lichtenstein, S. Lundström, N. Långström, A. Auton, K. M. Harris, G. W. Beecham, E. R. Martin, A. R. Sanders, J. R. B. Perry, B. M. Neale, B. P. Zietsch, Large-scale GWAS reveals insights into the genetic architecture of same-sex sexual behavior. Science 365 (2019).

106. B. P. Zietsch, M. J. Sidari, A. Abdellaoui, R. Maier, N. Långström, S. Guo, G. W. Beecham, E. R. Martin, A. R. Sanders, K. J. H. Verweij, Genomic evidence consistent with antagonistic pleiotropy may help explain the evolutionary maintenance of same-sex sexual behaviour in humans. Nat Hum Behav 5, 1251–1258 (2021).

107. UK Biobank Genetic Correlation Browser, Neale Lab (2018). https://ukbb-rg.hail.is.

108. C. Bycroft, C. Freeman, D. Petkova, G. Band, L. T. Elliott, K. Sharp, A. Motyer, D. Vukcevic, O. Delaneau, J. O’Connell, A. Cortes, S. Welsh, A. Young, M. Effingham, G. McVean, S. Leslie, N. Allen, P. Donnelly, J. Marchini, The UK Biobank resource with deep phenotyping and genomic data. Nature 562, 203–209 (2018).

109. A. D. Grotzinger, M. Rhemtulla, R. de Vlaming, S. J. Ritchie, T. T. Mallard, W. D. Hill, H. F. Ip, R. E. Marioni, A. M. McIntosh, I. J. Deary, P. D. Koellinger, K. P. Harden, M. G. Nivard, E. M. Tucker-Drob, Genomic structural equation modelling provides insights into the multivariate genetic architecture of complex traits. Nat Hum Behav 3, 513–525 (2019).

110. Sexual Offences Act (2003; https://www.legislation.gov.uk/ukpga/2003/42/contents).

111. S. Gil, Male victims of childhood sexual abuse by a male or female perpetrator. J. Trauma. Stress Disord. Treat. 03 (2014).

112. M. Mohler-Kuo, M. A. Landolt, T. Maier, U. Meidert, V. Schönbucher, U. Schnyder, Child sexual abuse revisited: a population-based cross-sectional study among Swiss adolescents. J. Adolesc. Health 54, 304–311.e1 (2014).

113. M. Ferragut, M. Ortiz-Tallo, M. J. Blanca, Victims and perpetrators of child sexual abuse: Abusive contact and penetration experiences. Int. J. Environ. Res. Public Health 18, 9593 (2021).

114. B. Mathews, D. Finkelhor, R. Pacella, J. G. Scott, D. J. Higgins, F. Meinck, H. E. Erskine, H. J. Thomas, D. Lawrence, E. Malacova, D. M. Haslam, D. Collin-Vézina, Child sexual abuse by different classes and types of perpetrator: Prevalence and trends from an Australian national survey. Child Abuse Negl. 147, 106562 (2024).

115. A. Panofsky, Misbehaving science: Controversy and the development of behavior genetics. 321 (2014).

116. J. Carlson, B. M. Henn, D. R. Al-Hindi, S. Ramachandran, Counter the weaponization of genetics research by extremists, Nature Publishing Group UK (2022). 10.1038/d41586-022-03252-z.

117. T. Caulfield, Spinning the Genome: Why Science Hype Matters. Perspect. Biol. Med. 61, 560–571 (2018).

118. H. Hopf, S. A. Matlin, G. Mehta, A. Krief, Blocking the Hype-Hypocrisy-Falsification-Fakery Pathway is Needed to Safeguard Science. Angew. Chem. Int. Ed Engl. 59, 2150–2154 (2020).

