## Supplementary for "Confounding Fuels Misinterpretation in Human Genetics"

John W. Benning[1,2]*, Jedidiah Carlson[3,4,5]*, Olivia S. Smith[3,4], Ruth G. Shaw[6], Arbel Harpak[3,4,+]
[1]Department of Ecology and Evolutionary Biology, Cornell University, Ithaca, NY, USA
[2]Department of Botany, University of Wyoming, Laramie, WY, USA
[3]Department of Integrative Biology, University of Texas at Austin, Austin, TX, USA
[4]Department of Population Health, University of Texas at Austin, Austin, TX, USA
[5]Department of Mathematics, Statistics, and Computer Science, Macalester College, St. Paul, MN, USA
[6]Department of Ecology, Evolution and Behavior, University of Minnesota, St. Paul, MN, USA

*these authors contributed equally

This PDF file includes:

Supporting Information for:
Confounding Fuels Misinterpretation in Human Genetics 1
- Supplementary Notes 3
  - Supplementary Note 1: Overview of the model used in Clark (2023) 3
  - Supplementary Note 2: Heritability and assortative mating estimates in other literature 5
  - Supplementary Note 3. Inconsistency of parameter estimates 7
  - Supplementary Note 4: Flawed tests for non-genetic influences 7
  - Supplementary Note 5: Construction of the occupational status index 9
  - Supplementary Note 6: Statistical artifacts influencing familial correlation estimates 10
    - a. Within-lineage correlations are highly attenuated 10
    - b. Inference about decay in correlations is substantially affected by pseudoreplication artifacts 12
    - c. Other potential sources of bias in familial correlations 13
  - Supplementary Note 7: Examining the utility of the persistence rate as a measure of social mobility 14
    - Familial correlations decreased over time 15
    - Sensitivities of the persistence rate parameter 16
  - Supplementary Note 8: Methods for reanalysis of Song & Zhang (2024) 16
- Supplementary Tables 19
  - Table S1. Parameter estimates reported in Clark (2023) Figure 1 and Table S2 are inconsistent with one another. 19
  - Table S2. Correlation matrix of individual status measures, including correlations with an individual's paternal wealth. 20

Table S3. Testing for equal mean status measures between fathers from surname lineages with <30 father-son pairs versus fathers from lineages with ≥30 father-son pairs (considering only unique individuals). 21
Table S4. Descriptive statistics of UK Biobank data used in our reanalysis of Song & Zhang (2024) 21
Table S5. Phenotypes with statistically significant genetic correlations with self-reported same-sex sexual behavior (SSB) in UK Biobank 23
Supplementary Figures 26
Figure S1. Exploring tests of environmental influences on social status in Clark (2023). 26
Figure S2. Familial correlations in occupational status vary widely across surname lineages. 27
Figure S3. Distributions of lineage-specific familial status correlations. 28
Figure S4. Variation and substructure among and within surname lineages. 29
Figure S5. Relationship between lineage-specific correlation in occupational status and lineage size, across relative types. 30
Figure S6. Distributions of familial status correlations stratified by sublineage. 31
Figure S7. Pseudoreplication of individuals in Clark (2023), illustrated here for the measure of occupational status (1780-1859). 32
Figure S8. Effect of pseudoreplication in Clark (2023) on variance and covariance estimates. 33
Figure S9. Empirical distributions of occupational status scores vary with genealogical distance and are distorted by pseudoreplication. 34
Figure S10. Mean and variance of birth year and status measures across genealogical distance. 35
Figure S11. The persistence parameter b is insensitive to the magnitude of familial correlations and describes only the rate at which they change as relatedness decreases. 36
Figure S12. Sensitivity of the persistence parameter, b, to changes in relative correlations. 37
Figure S13.Complementary analyses of temporal trends in familial correlations. 38
Figure S14. Path diagrams depicting results of Genomic SEM models with different postulated causal relationships between BSB, risk-taking, and number of children. 39
Figure S15. Partial genetic correlations between BSB and the measures considered in the alternative Genomic SEM models described in Fig. 3b. 40
Figure S16. Partial genetic correlations between number of children and the measures considered in the alternative Genomic SEM models described in Fig. 3b. 41
References 42

### Supplementary Notes

**Supplementary Note 1: Overview of the model used in Clark (2023)**

In this note, we elaborate on the math and assumptions of the inference model used in Clark (2023) (*1*). This model is based on earlier work by Gimelfarb (*2*), which synthesized the equations originally derived by Fisher (*3*) into a general linear model of genetic covariance between relatives under assortative mating. Under the assumptions outlined in (*2*), the genetic covariance between non-vertical relatives of degree $n$ (e.g., first cousins), $cov_{G,n}(X, Y)$, can be modeled as:

$$cov_{G,n}(X, Y) = h^2 (\frac{1+m}{2})^n \sigma_P^2 . \qquad \text{Eq. 1}$$

Here, $(X, Y)$ is a set of individuals and their relatives of degree $n$, $h^2$ is the narrow-sense heritability of the trait, $m$ is the genetic correlation between parents, and $\sigma_P^2$ is the phenotypic variance among individuals from whom $X$ and $Y$ are descended. (*1*) states that this is equivalent to the following model:

$$\rho_{P,n}(X, Y) = h^2 (\frac{1+m}{2})^n . \qquad \text{Eq. 2}$$

In this implementation,

$$\rho_{P,n}(X, Y) = cov_{P,n}(X, Y)/\sigma_{P,x}\sigma_{P,y} \qquad \text{Eq. 3}$$

is the phenotypic correlation between pairs of relatives $X$ and $Y$ of degree $n$, where $cov_{P,n}(X, Y)$ is the phenotypic covariance between relative pairs of degree $n$, $\sigma_{P,x}$ is the standard deviation of the trait among the set of records listed first in each relative pair, and $\sigma_{P,y}$ is that of the records listed second in each pair. We note that the implementation of Gimelfarb's model in (*1*) requires two additional assumptions: (i) the genetic and phenotypic covariance are equal ($cov_{G,n}(X, Y) = cov_{P,n}(X, Y)$), and (ii) $\sigma_{P,x} = \sigma_{P,y}$ for all values of *n* (i.e., the phenotypic variance,

$\sigma_P^2$, is constant across all genealogical relationships and over time). Taking the logarithm of both sides of Eq. 2 gives us:

$$log(\rho_{P,n}) = log(h^2) + n\,log(\frac{1+m}{2}).$$ Eq. 4 [equation 1 from (*1*)]

When $\rho_{P,n}(X, Y)$ measures the phenotypic correlation between vertical relationships (e.g., fathers and their sons), the equation includes an additional term that is a function of $r$, the phenotypic correlation between parents:

$$log(\rho_{P,n}) = log(h^2) + n\,log(\frac{1+m}{2}) + log(\frac{1+r}{2}).$$ Eq. 5 [equation 2 from (*1*)]

We note that (*1*) uses $\rho_n$ to represent the Pearson correlation coefficient, and not Spearman's rank correlation coefficient (which is customarily denoted as ρ); we follow this convention to differentiate from the phenotypic assortative mating parameter, $r$.

Combining these two equations, (*1*) fits the following linear regression model:

$$log(\rho_n) = a + n\,log(b) + c\,d_{lin}$$ Eq. 6 [equation 3 from (*1*)]

Where $d_{lin}$ is an indicator variable for single parent-child or single grandparent-grandchild relationships.

For clarity, we rewrite Eq. 6 as:

$$log(\rho_n) = \beta_0 + \beta_1 n + \beta_2 d_{lin} + \epsilon.$$ Eq. 7

Under this model, (*1*) obtains the following parameter estimates:

$$\widehat{h^2} = exp(\widehat{\beta_0})$$ Eq. 8

$$\hat{b} = exp(\widehat{\beta_1})$$ Eq. 9

$$\widehat{m} = 2\widehat{b} - 1 \qquad \text{Eq. 10}$$

A central assumption of Gimelfarb's (*2*) general linear model (Eq. 1) which extends to the model presented by Clark (*1*) (Eq. 2) is that the hereditary component of the quantitative trait/outcome being modeled is strictly determined by genotype. For sociobehavioral traits/outcomes in humans, however, we must consider some non-genetic transmissibility component, $c^2 > 0$ (following the notation of Cloninger et al. (*4*), who refer to this parameter as "cultural heritability;" we extend this definition to encompass material transmission and other non-genetic sources of similarity). Thus, assuming that non-genetic inheritance is at play for the socially-constructed traits considered in (*1*), $\widehat{h^2}$ in (*1*) is not an estimator of the narrow sense heritability, $h^2$, but of the total transmissibility:

$$t^2 = h^2 + c^2 + 2whc, \qquad \text{Eq. 11}$$

where $whc$ is the covariance between additive genetic and transmitted cultural factors within individuals [see equation 2 in (*4*), for example]. This partly explains unusually high estimates in (*1*), such as an estimate of $\widehat{h^2} = 0.94$ for occupational status (see Figure 1 from (*1*)).

Similarly, $\widehat{m}$ is described in (*1*) as an estimate of the genetic correlation between parents that are mating assortatively on some latent "social ability" phenotype. However, since socially constructed traits such as occupational status, higher education, house value, etc. have some non-genetic transmissible component ($c^2 > 0$), the true value of $m$ is a function of $t^2$ of the latent phenotype [as given by equation 4 of (*1*)], so it follows that the description of $m$ as a genetic parameter is mistaken. The parameter $m$ in (*1*) can therefore only be interpreted as the correlation between mates in a transmitted latent variable; *b* in turn can be interpreted as the log-additive decay of phenotypic correlations with degree of relatedness. Collado et al. (*5*) interpret *b* as representing "*how strongly* [socioeconomic] *advantages are transmitted from one generation to the next*." If we consider Fisher's model (*3*) as the generative process, this persistence of transmission is a result of assortative mating (regardless of whether the factor by which mates assort is fully, partly, or not at all genetic).

#### Supplementary Note 2: Heritability and assortative mating estimates in other literature

The suggestion in (*1*) that social inequality emanates from near-deterministic *genetic* inheritance of social status is at odds with recent literature (*5*–*8*). While estimates of narrow-sense heritability and the degree of assortative mating greatly depend on the population considered (geographically, genetically, socially and otherwise), definition of the trait, time, study characteristics and more, we discuss below some recent estimates of these parameters in genetic studies to contextualize estimates in (*1*).

Collado et al. (*5*) applied a model of genetic and cultural inheritance under assortative mating to a genealogical dataset from Sweden (similar in principle to that used in (*1*)) and estimated $m \approx 0.025$ for educational attainment (EA). Similarly, (*8*) estimated $m \approx 0.027$ for educational attainment using genomic data from ~400,000 residents of the United Kingdom.

The estimate of $m = 0.57$ in (*1*) is noted as "surprising" but is claimed to be in accord with two recent genomics studies of educational attainment, referring to (*9*, *10*)). First, (*9*) is cited in (*1*) as having found a "correlation [between mates] at trait-associated loci for educational attainment of 0.654," deemed compatible with $m = 0.57$. However, (*9*) states that 0.654 is the correlation between the observed phenotype of an individual and the phenotype that is predicted by their partner's genotypes at EA-associated SNPs. The authors of (*9*) acknowledge that this approach "cannot differentiate between direct assortment on a phenotype and assortment on a genetically correlated trait."

A more recent paper by Okbay et al. (*10*) reported the correlation in polygenic scores (PGS) for educational attainment between spouses to be 0.175. This study is cited in (*1*) with the assertion that "*since the polygenic index is a noisy measure of the full genetic educational potential*, *the full correlation will be significantly higher than this measured correlation*." There follows an attempt to derive what this "full correlation" should be for educational attainment by pivoting to the "analogous case of height" (it is unclear in what sense height should be considered analogous to educational attainment). Using Okbay et al.'s estimate of $r = 0.29$ as the observed phenotypic correlation in height between spouses and an assumed $h^2 = 0.8$ for height, (*1*) estimates that the "true" spousal genetic correlation ($m$) for height must be $rh^2 = 0.236$. Given that Okbay et al. estimated the spousal correlation in PGS for height to be 0.106, (*1*) reasons this is an underestimate of the true spousal genetic correlation (0.236) by a factor of 1.65 to 3.27. (*1*) then reasons that Okbay et al.'s spousal PGS correlation for EA (0.175) underestimates the "true" spousal genetic correlation by a similar degree, and concludes that "*the implied actual* [genetic] *correlation averages 0.39* $[0.175 \times \frac{0.236}{0.106}]$, *with a 95% CI* [confidence interval] *of 0.29 to 0.57…so, the 0.175 [PGS] correlation observed between partners for educational attainment is potentially consistent with a true genetic correlation of 0.57*."

This numerological reasoning has no basis in quantitative genetic theory. First, we note that this claim in (*1*) relies on strong and unfounded assumptions about (i) the equivalency of the genetic architectures of two complex anthropometric and behavioral traits and (ii) the extent to which polygenic scores (and the GWAS associations they are constructed from) are free from bias and capture underlying genetic effects for each of these traits. Second, we point out that the 95% confidence interval given for the implied spousal genetic correlation point estimate—[0.29, 0.57]—is not derived from the probability distribution of the parameter of interest (i.e. the spousal PGS correlation for educational attainment), so this CI cannot be interpreted as a measure of statistical uncertainty. While a deeper investigation of the relationship between spousal PGS correlations and $m$ (and how to appropriately quantify uncertainty) is beyond the scope of our reanalysis, we emphasize that (*1*) fails to reconcile its estimate of $m = 0.57$ with other studies (namely, (*5*) and (*8*)) that obtained estimates of $m$ more than 20 times lower using methods that rigorously attempt to control for confounding.

**Supplementary Note 3. Inconsistency of parameter estimates**

The model summary statistics ($b$ and $h^2$ for 9 different traits) presented in Figure 1 of (*1*) are claimed to be the same as those found in Supplementary Table S2 of (*1*), but these values are different (for example, the heritability of occupational status among men born 1780-1859 appears to be ~0.90-0.95 in Figure 1, but is given as 0.72 in Table S2). Percent differences in estimated trait heritability values between Table S2 and Figure 1 range from -27% to 72%. Our attempts to reproduce the results in (*1*) (using a weighted least-squares regression as described in (*1*), where the weights for each $\rho_n$ are equal to $1/\sigma_n$) tend to agree more closely with values in Figure 1 of (*1*), suggesting that the parameter estimates in Table S2 of (*1*) may have been derived from a different model or used a different version of the dataset (**Table S1**).

**Supplementary Note 4: Flawed tests for non-genetic influences**

The main analyses in (*1*) assume all transmissibility is due to narrow-sense heritability. Then, (*1*) uses two analyses to rule out "environmental influences" on social status post-hoc.

First, it is shown that the father-son correlations in occupational status and education are relatively unaffected by the son's age at his father's death [(*1*) Figure 4]. The high correlation between father and son measure values even when the father is largely absent from the son's life is taken in (*1*) to indicate the effect of a father on the social status of their child being largely limited to genetic heritability. But many pertinent non-genetic factors and environmental conditions can be transmitted regardless of a father's presence, such as familial wealth, place of residence, interactions with father's family, and familial traditions of occupation. Assortative mating is also likely to buffer any decrease in paternal environmental effects owing to a father's death, as mothers tend to be correlated with fathers in attributes (literacy, etc.) that shape child outcomes.

We illustrate the predictiveness of non-genetic paternal effects in the absence of fathers by examining whether the association between paternal wealth and offspring status changes when fathers die early in a son's life. In these data, there is no significant effect of a son's age at father's death on the correlation between paternal wealth and son's educational attainment or occupational status, even when the father dies early in his son's life (**Fig. S1a**). This accords with the father-son correlations in education and occupational status that are also high for fatherless boys, as paternal wealth is strongly associated with these status measures (**Fig. 2**, **Table S2**). The absent/present father analysis in (*1*) therefore does not help disentangle genetic from non-genetic transmission or speak to the relative influence of environmental factors on variation in status.

Second, status measures were shown to be transmitted equally through maternal and paternal lines, whereas wealth was transmitted more strongly through paternal lines [(*1*) Figure 3]. It is unclear why similar statistical associations of social status with mothers and fathers is taken as evidence for genetic underpinnings (here, maternal and paternal effects are proxied via grandparental measures). Such a pattern is indeed consistent with genetic transmission, but could also be consistent with many forms and patterns of non-genetic transmission. For instance, there is no *a priori* reason to think that non-genetic transmission of literacy from mothers to offspring would, on average across a population, be stronger or weaker than transmission by fathers. Though of course the true parental effects are unlikely to be exactly equal given non-genetic transmission, the power to accurately and precisely estimate these effects, and thus rule out their inequality, is limited in most data sets. Furthermore, in (*1*) the models for higher education, occupational status, and wealth are fit to different data sets with different distributions of wealth and status (**Fig. S1b**), such that a comparison of coefficients across models is uninformative. We performed the same analysis on the subset of (*1*)'s data that had complete information (higher education, occupational status, and wealth) for each individual (N = 817). Using this analysis that rules out differences due to underlying data sets, the maternal wealth and paternal wealth effects on an individual's own wealth are statistically indistinguishable (**Fig. S1c**).

**Supplementary Note 5: Construction of the occupational status index**

Beyond model misinterpretation, a source of inflation in $t^2$ and $b$ in (*1*) may be upstream choices in the collection and preparation of the data. For example, the occupational status index used in (*1*) was devised by the paper's author and others in a recent preprint (*11*). The data underlying the index are 1.6 million marriage records in England across years 1837-1939, which include data on occupations for brides, grooms, and both of their fathers. (*11*) condense the more than 100,000 occupation description strings in those data into 442 occupational categories (listed in the Appendix Table A.3 in (*11*)). For comparison, a standard occupational status index for this period, the HISCAM-GB, uses 1,300 occupational categories (*11*, *12*). After individuals were assigned an occupation, Goodman's RCII association model (*13*) was used to generate a status index for each occupation. Though the specific details of model specification are not presented in (*11*), RCII models generally proceed as follows. The core idea in such analyses is that social stratification will be reflected in patterns of occupational interactions. The analysis takes as input a contingency table, where rows represent one individual's occupation, and columns represent the paired individual's occupation, with cell counts corresponding to the number of cases where this combination of occupations is observed. The RCII approach then fits a log-multiplicative model to these categorical data, of the general form

$$log(F_{ij}) = \mu + \mu_i^R + \mu_j^C + \beta\phi_i\varphi_j, \qquad \text{Eq. 12}$$

where $F_{ij}$ is the expected cell frequency, $\mu$ is the main effect, $\mu^R$ is the row effect, $\mu^C$ is the column effect, $\beta$ is the association parameter measuring the association between row and column variables, and $\phi_i$ and $\varphi_j$ are the unknown row and column scores (indices) to be estimated (*14*). When used to estimate status indices, row and column scores of a given occupation are constrained to be equal. An algorithmic procedure that iteratively assigns scores to occupational categories is used to estimate the occupation index scores that maximize the fit between the observed counts ($f_{ij}$) in each cell and the model-predicted counts ($F_{ij}$). Thus, this procedure maximizes the correlation in occupational status index for whatever pair of individuals is analyzed (bride-groom, father-son, etc.). In (*11*), index estimation was performed according to this approach separately for the father/son occupation associations and for the father-in-law/son-in-law associations; the average of these indices was used as the overall index of occupational status in (*1*). Specifically for the data collection and preparation choices (e.g. choice and coding scheme for occupational categories) of (*11*), this method resulted in father/son occupational status correlations ~30% higher than those based on other widely used indices of occupational status (*11*). This methodological choice may be, at least in part, contributing to the unusually high correlations of relatives at the heart of the arguments in (*1*).

**Supplementary Note 6: Statistical artifacts influencing familial correlation estimates**

The conclusions of (*1*) are based on the observation that estimates of familial correlation for measures of social status decay linearly with genealogical distance, regardless of the status measure or time period considered. Our reanalysis of the data reveals that these correlation estimates–and the manner in which they change with genealogical distance–are heavily affected by statistical artifacts. First, we show that the overall correlation patterns reported in (*1*) do not consistently reflect within-lineage correlation patterns (**Note 6a; Figs. S2-S6**). Next, we show that in (*1*)'s analyses, all pairs of relatives are treated as mutually independent observations, yet many individuals are represented in multiple relative pairs from which the correlation coefficients are estimated (**Note 6b; Figs S7-S9**). For example, the (1780-1859) occupational status correlation for fourth cousins is calculated from 17,382 pairs, derived from only 1,878 unique individuals from just 31 surname lineages. This treatment of non-independent data points as mutually independent is commonly known as pseudoreplication, with a known effect of driving underestimates of statistical uncertainty (*15*). Here, however, the pseudoreplication is non-uniform: as genealogical distance increases, individuals are increasingly re-counted in more relative pairs, but these individuals are from a diminishingly smaller selection of lineages (**Fig. S7**, **Fig. S8; Fig. S9**). Because of the non-uniform pseudoreplication, the point estimates of familial correlations in (*1*) may even be biased. Indeed, when we avoided pseudoreplication by randomly sampling pairs of relatives (one pair from each surname lineage), point estimates of sample correlations do not even decrease monotonically with genealogical distance; in addition, their noisiness prevents making confident assertions about the persistence of these correlations with genealogical distance (**Fig. 2b**). (We note that adjusting for pseudoreplication in this way is certainly not a complete solution to the problems of pseudoreplication and familial structure, could introduce its own biases, and is only one of many possible methods of adjustment for these problems.) Finally, we show that individuals comprising more distant relationship types are also wealthier, born more recently, and have higher occupational status (**Note 6c; Figs. S9-S10**). This trend may be partially due to sampling biases or temporal change in the population distribution of occupational status that are unaccounted for (**Fig. S10**). Together, the core flaws of model misspecification and statistical artifacts call into question inferences drawn in (*1*) based on familial correlations in status.

##### a. Within-lineage correlations are highly attenuated

The core results in (*1*) are based on "lineage-agnostic" correlations in social status, which are calculated between individuals in all pairs of a given genealogical relationship, ignoring the surname lineage to which each pair belongs and ignoring heterogeneity among and within these lineages. We find that within-surname distributions of correlations are highly variable across surnames. This is true for all 11 genealogical relationships and for all nine measures of status (**Figs. S2a, S3**). The central tendencies of these surname-specific correlations are substantially lower than the corresponding lineage-agnostic correlations reported in Table 2 of (*1*). Beyond the attenuation of correlations that is to be expected when conditioning on a given family, there is remarkable heterogeneity among lineages: the medians and modal values of the surname-specific correlations are near zero for all genealogical relationships beyond first cousins (**Figs. S2a, S3**), but some surnames show high correlations, even among distant cousins (**Figs. S2, S3, S4a**).

This finding may also be partly explained by the transmissibility of social status varying among families, reminiscent of the Scarr-Rowe effect, where the estimated heritability of cognitive ability varies by socioeconomic status (*16*–*21*), and is consistent with arguments that when families vary in their access to, use of, and transmission of cultural tools, resources, and norms, the transmissibility of phenotypes impacted by these non-genetic sources of variation will also vary among families (*22*, *23*).

An illustrative example is surname lineage number 1436 (**Fig. S4b**). Applying the regression model of (*1*) to the occupational status correlations calculated within this surname lineage among men born 1780-1859, we obtain an estimate of $b = 0.99$—there is no decay in correlation, even out to fourth cousin pairs. The reason for this result is strong structure in this surname lineage. For instance, the fourth cousin pairs come from four different sublineages (i.e., descended from a different great-great-great grandfather, as we have identified using the *pidf* variable in the data). The correlation within each sublineage is 0, but because sublineages differ in the average occupational status of their members, the sublineage-agnostic correlation among all fourth cousin pairs with this surname is 0.63 (**Fig. S4b**). Importantly, similar substructure is what drives the lineage-agnostic correlation when fourth cousin data from multiple surnames are combined (**Fig. S4c**).

For first-degree relatives, within-surname heterogeneity tends to be more pervasive in surnames with more observed relative pairs, resulting in surname-specific correlations that tend to be dramatically higher (*1*) (**Fig. S2b-c**, **Fig. S5**). As genealogical distance increases, however, the relationship between the number of surname pairs and surname-specific correlation gradually dissipates (**Fig. S5**). At a more granular level, when we condition on sublineages, the correlations (and thus the covariances) are almost always near 0, across all genealogical relationships (**Fig. S6**).

This trend can also be illustrated through the dependency of within-lineage correlations on sample size among first-degree relatives: surnames with more represented members (≥30 father-son or full sibling pairs) are highly correlated (median $r = 0.46$ for full siblings and $r = 0.52$ for father-son pairs) and surnames with fewer represented members (<30 father-son or full sibling pairs) are less correlated (median $r = 0.01$ for full siblings and $r = 0.18$ for father-son pairs). These differences cannot be attributed to differences in estimation noise due to varying sample sizes.

##### b. Inference about decay in correlations is substantially affected by pseudoreplication artifacts

The dataset used in (*1*) consists of all observed pairs of a given relationship, and each individual will typically be represented in multiple records. For example, one family of ten brothers (all sons of individual 80207) is represented by $\binom{10}{2} = 45$ sibling pairs in the data, all of which are treated as independent observations. This means that even though the pairwise records are all distinct from one another, when data from the same individual is repeated in multiple pairwise records, these records are considered "pseudoreplicates" of one another, because they are not mutually independent. Pseudoreplication can lead to a multitude of problematic statistical artifacts and inference errors, some of which are documented in (*24*) (see (*25*) for a related discussion of how genealogical pseudoreplication impacts inference in the context of genetic association studies). In (*1*), the effects of pseudoreplication are increasingly pervasive for more distant relationships, where there are generally exponentially more pairwise relationships within a given lineage (**Fig. S7**).

Pseudoreplication among relative pairs can affect the lineage-agnostic correlation estimates in two ways: by altering the covariance between relatives and by altering the sample variance. For close relationships (out to first cousins), pseudoreplication causes both the covariance and variance of occupational status to increase, but for more distant relatives, the covariance and variance of occupational status decrease (**Fig. S8**). Though the sample variances change at most by ~20%, the changes in covariance can be dramatic: for third cousins once removed, the covariance decreases nearly 80% when pseudoreplicated records are included. To mitigate this effect of pseudoreplication on relative correlation estimates, we sampled a single pair of relatives of each relationship type from each surname. While mitigating pseudoreplication, this down-sampling reduces the sample size significantly. We therefore performed 1,000 random bootstrap samples and took the averages of the relative correlations as an estimate. Note that this approach also removes some higher-order pseudoreplication effects; for example, suppose we have a pair of brothers who are fourth cousins with another pair of brothers, producing four pairwise fourth cousin relationships. Even though we can downsample these into two pairs that contain mutually exclusive individuals (removing the first-order effects of pseudoreplication), each pair of brothers share the same father, which will presumably have a strong influence on their status, so there is some degree of second-order pseudoreplication caused by resemblance between siblings. This sampling strategy does not account for other forms of pseudoreplication (e.g. the same individual being represented in multiple pairs across different relationship types or relatedness via a recent maternal common ancestor), and we are unable to resolve potential biases that might arise due to genealogical or social relationships across different surname lineages in these data. As shown in **Fig. 2b**, pseudoreplication affected lineage-agnostic occupational status correlations estimated by (*1*). When applied to the revised correlation estimates, the log-linear regression model of (*1*) explains only 29% of the variation in relative correlations.

##### c. Other potential sources of bias in familial correlations

The distributions of each status measure differ across the subsets of different genealogical relationships, indicating these subsets do not represent the same cross-sections of the population (**Fig. S9**). For instance, among men in the dataset born 1780-1859, the mean occupational status among unique individuals represented in father-son pairs (37.5) is ~10% lower than among members of fourth cousin pairs (mean occupational status = 41.6; $p = 3.58 \times 10^{-23}$) (**Fig. S10**). As genealogical distance increases, the individuals comprising each subset also tend to be wealthier, born more recently, and have greater variance in occupational status (**Fig. S10**). One potential explanation for these differences in sample distributions is that social or demographic changes have altered the true population distributions of status measures over this time period. Specifically, temporal changes to the population variance of occupational status are important to consider: because the denominator of the familial correlations modeled by (*1*) is assumed to be an estimator of the population variance [see (*2*)], and because more distant relatives in the data tend to have been born more recently, the changes in correlation across genealogical distance may partly reflect changes in the population distribution over time.

Another partial driver of the differences in familial correlation may be sampling and ascertainment biases that render the data unrepresentative of the broader sample/population. The selection criteria for lineages to be included in the dataset, described in Appendix 01 of (*1*), acknowledges that this dataset is largely a convenience sample, stating: "*lineages were chosen for inclusion based on their completeness, and either the public posting of the lineages or their creators' willingness to share the data*," but the potential biases that might be induced by these particular selection criteria are not addressed. In addition, we find that the average wealth and occupational status are significantly higher (and the year of birth significantly earlier) for fathers from large lineages compared to those of small lineages (**Table S3**).

**Supplementary Note 7: Examining the utility of the persistence rate as a measure of social mobility**

In (*1*), the "persistence rate", $b$, measures the rate of decay in familial correlations with increasing genealogical distance—i.e., if the correlation in occupational status between first cousins is 0.8 that of uncles and nephews, which is 0.8 that of full siblings and so on, then $b = 0.8$. In (*1*), the large estimated value of the persistence rate, $b \approx 0.79$, and its stability across social status measures and between two time periods (for two of the measures), are taken as evidence for a persistence of social status and rate of social mobility that has been largely unaffected by societal changes:

> "*People in 2022 remain correlated in outcomes with their lineage relatives in exactly the same way as in preindustrial England.*"

> "*The vast social changes in England since the Industrial Revolution, including mass public schooling, have not increased, in any way, underlying rates of social mobility*".

Across time-based comparisons presented in the paper and all degrees of relationship [(*1*) Table 2], 16/22 correlations decrease between the two time periods analyzed (on average, decreasing 31%). How could the estimate of $b$ lead to conclusions that contrast what is suggested by the vast majority of data points from which it was inferred, and contradict common measures of mobility so starkly? Below we describe three aspects of $b$ that bring into question this parameter's suitability as an informative measure of social mobility.

First, even when familial correlations do not decay monotonically with degree of relatedness, a large estimate of $b$ is likely. For example, fitting the regression model of (*1*) to the correlation estimates adjusted for pseudoreplication shown in **Fig. 2b** (where there is no obvious relationship and the linear fit has $R^2 = 0.29$) yields an estimate of $b = 0.93$.

Second, even with a strong linear fit to the data, $b$ is uninformative as to the magnitude of familial correlations and describes only the rate of decrease in the correlation as relatedness declines. For illustration, $b$ approaches 1 as the association between familial correlation and relatedness nears zero [(*1*) Eq. 3]. Importantly, familial correlations in a status measure could systematically increase or decrease by any amount, with $b$ remaining unchanged (**Fig. S11**).

Third, the estimate of $b$ can be greatly affected by invalid statistical modeling. We highlight two examples. First, (*1*) effectively standardizes trait variance across different relationship types. This procedure obscures systematic trends that we observed in trait mean and variance across these subsets, which are incompatible with (*3*) and (*2*) model assumptions (**Supplementary Note 1**; **Supplementary Note 6c**; **Figs. S8, S9, S10**). A second example lies in increased sensitivity of the estimate of *b* to small changes in the correlations of distant relatives, but relative insensitivity to the correlations of closer relatives ( see “Sensitivities of the persistence rate parameter” below; **Fig. S12**). As we have seen empirically, estimates of status correlations for distant relatives were also most sensitive to pseudoreplication and substructure among and within lineages due to sampling bias, noisiness, and/or temporal changes in the population distribution (**Supplementary Note 6**; **Figs. 2b**, **S8**, **S9**, **S10**).

##### Familial correlations decreased over time

Given these characteristics of $b$, it is worth considering more established metrics of social mobility, such as changes in parent-offspring correlations in social status over time (*26*–*28*), in data from (*1*). As we show in the main text, parent-offspring correlations in occupational status, higher education, and literacy (the only three status measures with data prior to the 20th century), as well as wealth, decrease over time (**Figs. 2c**, **S13**). Correlations in occupational status and higher education for sibling, grandparent/grandchild, uncle/nephew, and first cousin relationships also tend to decrease (**Fig. S13**).

##### Sensitivities of the persistence rate parameter

We used simulations to understand how estimates of the persistence rate parameter, $b$, vary across a wide range of possible patterns of familial correlations (from full sibling to fourth cousins). We did not include direct descendant relationships (parent-child, grandparent-grandchild) for simplicity. For each instance of the simulation, we generated correlations for each relative type as follows: for full siblings, a correlation was drawn from $U(0, 0.7)$. For 4th cousins, a correlation was drawn from $U(0, 0.1)$, with the constraint that its value was less than that of the full sibling correlation. We then set *b* as the exponential of the slope describing the relationship between the logarithms of those two correlations:

$$log(b) = \frac{log(\rho_{4th\ cousin}) - log(\rho_{full\ sibling})}{8}$$

We then set correlations for the remaining relationships (from siblings once removed to 3rd cousins once removed) as

$$log(\rho_n) = log(h^2) + n\, log(b) \qquad \text{[Equation 1 from (}\textit{1}\text{)]}$$

where $n$ is the degree of relatedness (i.e., $n = 1$ for full siblings and $n = 9$ for fourth cousins; **Supplementary Note 1**). Each correlation was then adjusted by random deviate, drawn from a normal distribution with zero mean and a standard deviation of 0.2. Values were exponentiated to return to the original scale and then bounded between 0 and 1 to ensure validity. Using the set of generated correlations, the same linear regression model described above was fitted, and $b$ was estimated. The simulation results (**Fig. S12**) show that $b$ is most sensitive to distant relative correlations and is nearly guaranteed to exceed 0.7 for cases having correlation between 4th cousins exceeding 0.04.

**Supplementary Note 8: Methods for reanalysis of Song & Zhang (2024)**

Here we describe our methods for reproducing and expanding the analysis of (*29*). Using Version 3 of the imputed genetic data for 500k participants from the UK Biobank (*30*) (along with phenotype information and metadata for this sample, released on 2024-01-19) we followed the paper's published methods for data preprocessing, defining measures of sexual behavior and reproductive success, performing GWASs for these measures (and self-reported risk-taking), and modeling the resulting genetic correlations using Genomic Structural Equation Modeling (Genomic SEM) (*31*). The authors of (*29*) provided us with code to execute their GWAS pipeline using the PLINK (*32*) and REGENIE software (*33*), and GenomicSEM code was published in the paper's supplementary material.

Descriptive statistics comparing Song and Zhang's final dataset with our closest reproduction of their analysis are shown in **Table S4**. Note slight differences in our resultant post-filtering sample size and number of variants tested for association, which we attribute in part to unknown differences in (i) additional participants who withdrew their consent to include data in the UK Biobank since (*29*) conducted their analysis, (ii) the implementation of the k-means clustering algorithm used to screen samples based on genetic ancestry, and (iii) the order of operations used to filter samples and variants. Also note that we only considered autosomal variants in our reanalysis, whereas (*29*) included variants on the X chromosome. Despite these differences, the raw Pearson correlation coefficient between allele effect sizes from Song and Zhang's published GWAS summary statistics for BSB in males and those from our reproduction attempt was 0.941, and the LD Score Regression (*34*) genetic correlation between the two was 0.999. This suggests that any discrepancies between Song and Zhang's GWAS summary statistics and those from our reproduction attempt had a negligible impact on the estimation of genetic correlations and results of Genomic SEM (compare Fig 2b in (*29*) with **Fig. S14a** here). For consistency across our analyses, in particular the analysis of **Fig. 3b**, we used the summary statistics from our GWASs for these measures, rather than those published by (*29*).

We investigated whether the Genomic SEM model presented in (*29*) could be distinguished from models with alternative postulated causal relationships among BSB in males, risk-taking in males, and number of children (**Fig. S14b-d**). We also note that (*29*) misrepresented the causal paths in their model as horizontal pleiotropy between BSB and risk-taking (i.e., the same genes affect both phenotypes through independent biological pathways), when they actually modeled and graphically portrayed this relationship as vertical pleiotropy (alleles affect risk-taking behavior, which is a causal risk factor for BSB, but no alleles affect BSB independently from this pathway). Notably, when there are only three measures considered, Genomic SEM is incapable of providing statistical support for different modes of pleiotropy or particular causal pathways (*31*)—if we simply reverse the postulated causal path between risk-taking and BSB in the model, we obtain identical partial genetic correlation estimates.

We then investigated whether measures other than risk-taking could explain the abolishment of genetic correlation between BSB in males and number of children. We performed GWASs for a selection of other measures in the UK Biobank sample (shown in **Fig. 3b** and **Fig. S15**), using the same GWAS pipeline that (*29*) used for their focal phenotypes. Each of these measures were selected based on prior evidence of having a significant genetic correlation with risk-taking behavior, same-sex sexual behavior, and/or number of children (e.g. **Supplementary Table 5**). For each of these phenotypes/measures, we implemented a Genomic SEM (*31*) model with the same postulated causal structure as Song and Zhang's model, but replaced risk-taking behavior with that alternative phenotype/measure. We evaluated these models on the basis of whether adjusting for a given measure resulted in a partial genetic correlation between BSB in males and number of children that was not statistically different from zero.

### Supplementary Tables

**Table S1. Parameter estimates reported in Clark (2023) Figure 1 and Table S2 are inconsistent with one another.**
Attempts to reproduce these results are consistent with (1) Figure 1, but not (1) Table S2. In (1) $t^2$ is referred to as $h^2$.

| | Clark (2023) Figure 1 | | Clark (2023) Table S2 | | WLS from raw data | | OLS from correlations in Clark (2023) Table 2 | |
|---|---|---|---|---|---|---|---|---|
| | $t^2$ | b | $t^2$ | b | $t^2$ | b | $t^2$ | b |
| Modern Status | 0.39 | *0.81* | 0.47 | *0.77* | 0.39 | *0.80* | 0.39 | *0.81* |
| log(House Value) | 0.36 | *0.82* | 0.41 | *0.80* | 0.37 | *0.81* | 0.36 | *0.82* |
| Company Director | 0.11 | *0.82* | 0.19 | *0.78* | 0.13 | *0.86* | 0.11 | *0.82* |
| Index Mult. Deprivation | 0.36 | *0.72* | 0.36 | *0.73* | 0.39 | *0.70* | 0.36 | *0.72* |
| Literacy | 0.41 | *0.84* | 0.43 | *0.85* | 0.47 | *0.82* | 0.47 | *0.83* |
| Occ. Status (1780-1859) | 0.94 | *0.75* | 0.72 | *0.81* | 0.93 | *0.75* | 0.94 | *0.75* |
| Occ.Status (1860-1919) | 0.66 | *0.8* | 0.65 | *0.79* | 0.66 | *0.80* | 0.66 | *0.80* |
| Higher Ed (1780-1859) | 0.57 | *0.72* | 0.63 | *0.77* | 0.36 | *0.83* | 0.58 | *0.76* |
| Higher Ed (1860-1919) | 0.58 | *0.76* | 0.42 | *0.77* | 0.46 | *0.75* | 0.58 | *0.71* |

**Table S2. Correlation matrix of individual status measures, including correlations with an individual's paternal wealth.**

N/A values indicate the absence of individuals in the data set with values recorded for both measures.

| | Higher education | Occupational status | Index Mult. Deprivation | log(House Value) | Modern status | Literacy |
|---|---|---|---|---|---|---|
| Higher education | | | | | | |
| Occupational status | 0.59 | | | | | |
| Index multiple deprivation | 0.11 | 0.24 | | | | |
| Log (house value) | 0.26 | 0.45 | 0.54 | | | |
| Modern status | 0.25 | 0.47 | 0.81 | 0.85 | | |
| Literacy | 0.09 | 0.31 | *N/A* | *N/A* | *N/A* | |
| **Paternal wealth** | **0.48** | **0.66** | **0.19** | **0.36** | **0.35** | **0.26** |

**Table S3. Testing for equal mean status measures between fathers from surname lineages with <30 father-son pairs versus fathers from lineages with ≥30 father-son pairs (considering only unique individuals).**

| Variable | Small sample group (<30 father-son pairs) | | Large sample group (≥30 father-son pairs) | | T-test p-value |
|---|---|---|---|---|---|
| | Mean | Standard deviation | Mean | Standard deviation | |
| Occupational status | 35.4 | 12.1 | 37.9 | 16.0 | 1.66e-6 |
| Wealth | -3.61 | 2.06 | -2.50 | 2.65 | 5.34e-5 |
| Birth year | 1807 | 17.8 | 1803 | 20.4 | 3.88e-6 |

**Table S4. Descriptive statistics of UK Biobank data used in our reanalysis of Song & Zhang (2024)**

| Dataset | Sample size | Number of variants tested in GWAS |
|---|---|---|
| Song & Zhang 2024 | 452,557 | 9,371,426 (autosomes + chrX) |
| Our reproduction | 459,676 | 9,335,673 (autosomes only) |

**Table S5. Phenotypes with statistically significant genetic correlations with self-reported same-sex sexual behavior (SSB) in UK Biobank**
(white British male sample only; data obtained from https://ukbb-rg.hail.is/rg_browser/)

| Phenotype | Genetic correlation with SSB | SE | p-value |
|---|---|---|---|
| Manifestations of mania or irritability: I was more talkative than usual | 0.7386 | 0.2425 | 0.002321 |
| Workplace very hot: Often | 0.6553 | 0.1977 | 0.000917 |
| Ever taken cannabis | 0.6382 | 0.1146 | 2.55E-08 |
| Felt hated by family member as a child | 0.6109 | 0.2233 | 0.006216 |
| Belittlement by partner or ex-partner as an adult | 0.5984 | 0.221 | 0.006772 |
| Ever contemplated self-harm | 0.5979 | 0.224 | 0.007617 |
| Tinnitus: Yes, now most or all of the time | 0.5614 | 0.1769 | 0.001503 |
| Lifetime number of sexual partners | 0.5228 | 0.07805 | 2.10E-11 |
| Victim of physically violent crime | 0.5152 | 0.193 | 0.007594 |
| Particulate matter air pollution (pm10); 2007 | 0.4856 | 0.1499 | 0.0012 |
| Townsend deprivation index at recruitment | 0.4594 | 0.106 | 1.46E-05 |
| Nitrogen dioxide air pollution; 2007 | 0.4503 | 0.1304 | 0.000553 |
| Nitrogen dioxide air pollution; 2005 | 0.4454 | 0.133 | 0.00081 |
| Nitrogen dioxide air pollution; 2006 | 0.4362 | 0.1285 | 0.000688 |
| Ever smoked | 0.4164 | 0.08514 | 1.01E-06 |
| Particulate matter air pollution (pm2.5); 2010 | 0.4144 | 0.1403 | 0.003143 |
| Illness, injury, bereavement, stress in last 2 years: Financial difficulties | 0.4134 | 0.1159 | 0.000363 |
| Loneliness, isolation | 0.4094 | 0.1062 | 0.000116 |

| | | | |
|---|---|---|---|
| Ever unenthusiastic/disinterested for a whole week | 0.4057 | 0.1412 | 0.004059 |
| Nitrogen dioxide air pollution; 2010 | 0.393 | 0.1323 | 0.002964 |
| Ever thought that life not worth living | 0.3783 | 0.1407 | 0.007166 |
| Workplace very noisy: Often | 0.37 | 0.1423 | 0.009321 |
| General happiness | 0.3663 | 0.1152 | 0.001474 |
| Tobacco smoking: Ex-smoker | 0.3639 | 0.1172 | 0.001902 |
| Smoking status: Previous | 0.3395 | 0.08287 | 4.18E-05 |
| Nitrogen oxides air pollution; 2010 | 0.3324 | 0.1262 | 0.008424 |
| Seen a psychiatrist for nerves, anxiety, tension or depression | 0.326 | 0.1137 | 0.004149 |
| Smoking status: Current | 0.3256 | 0.0933 | 0.000483 |
| Current tobacco smoking | 0.3229 | 0.09214 | 0.000458 |
| Risk taking | 0.3201 | 0.08795 | 0.000273 |
| Coffee type: Ground coffee (include espresso, filter etc) | 0.2826 | 0.08984 | 0.001657 |
| Seen doctor (GP) for nerves, anxiety, tension or depression | 0.2759 | 0.09117 | 0.002476 |
| Lamb/mutton intake | 0.2637 | 0.09848 | 0.007412 |
| Irritability | 0.2347 | 0.08559 | 0.006104 |
| Corneal resistance factor (left) | 0.216 | 0.08028 | 0.007122 |
| Wheeze or whistling in the chest in last year | 0.2114 | 0.08114 | 0.00919 |
| Age first had sexual intercourse | -0.2331 | 0.06554 | 0.000375 |
| Number of vehicles in household | -0.2866 | 0.1019 | 0.004913 |
| Length of time at current address | -0.2875 | 0.1066 | 0.006963 |
| Coffee type: Instant coffee | -0.3015 | 0.0976 | 0.002004 |
| Reason for reducing amount of alcohol drunk: Other reason | -0.3409 | 0.1264 | 0.006987 |

| | | | |
|---|---|---|---|
| Past tobacco smoking | -0.3711 | 0.07559 | 9.16E-07 |
| Smoking status: Never | -0.3784 | 0.07401 | 3.16E-07 |
| Felt loved as a child | -0.3939 | 0.1328 | 0.003013 |
| How are people in household related to participant: Husband, wife or partner | -0.3939 | 0.107 | 0.000231 |
| Illness, injury, bereavement, stress in last 2 years: None of the above | -0.4063 | 0.1357 | 0.002762 |
| Bipolar and major depression status: No Bipolar or Depression | -0.4861 | 0.1483 | 0.001044 |
| Type of accommodation lived in: A house or bungalow | -0.4937 | 0.135 | 0.000255 |
| Types of transport used (excluding work): Car/motor vehicle | -0.5126 | 0.1268 | 5.28E-05 |
| Number in household | -0.5864 | 0.1544 | 0.000145 |

### Supplementary Figures

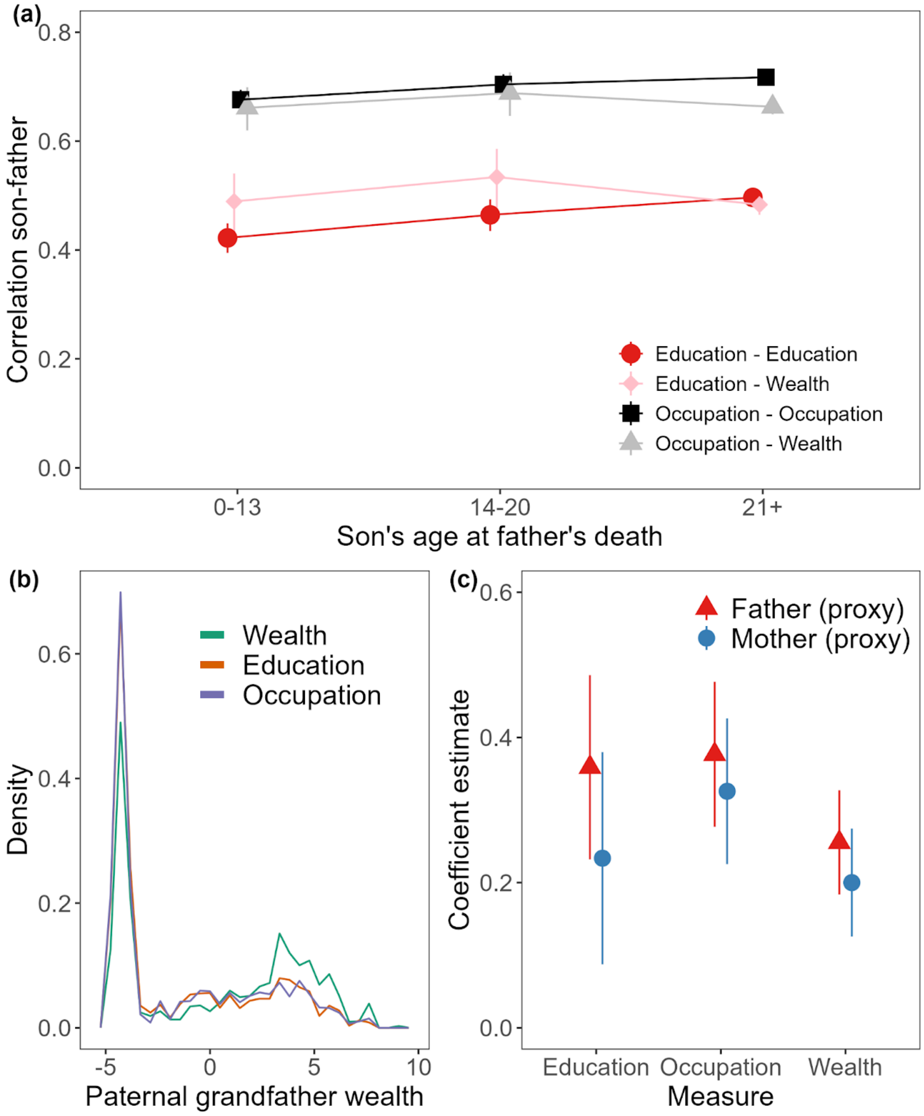


**Figure S1. Exploring tests of environmental influences on social status in Clark (2023).** **(a)** Father-son correlations (95% CI) in education (red), occupation (black), son education with father wealth (pink), and son occupation with father wealth (grey), across different bins of the son's age at his father's death. Compare to Figure 4 in (*1*). **(b,c)** maternal and paternal effects on offspring status measures and wealth are statistically indistinguishable when data used across these three regressions are identical (Supplementary Note 4). Compare to Figure 3 in (*1*). **(b)** shows distributions (30 bins) of log(paternal grandfather wealth) for individuals comprising each data set in the original analysis of (*1*); models of wealth were fit to a data set with more high wealth individuals. In **(c)**, coefficients (95% CI, clustered on fathers as in (*1*)) are shown for linear regressions of 'grandchild measure ~ maternal grandfather measure + paternal grandfather measure', using a data set of complete records. Here and in the analyses of (*1*), maternal and paternal grandfather measures are used as proxies for maternal and paternal measures, due to no information on mothers directly.

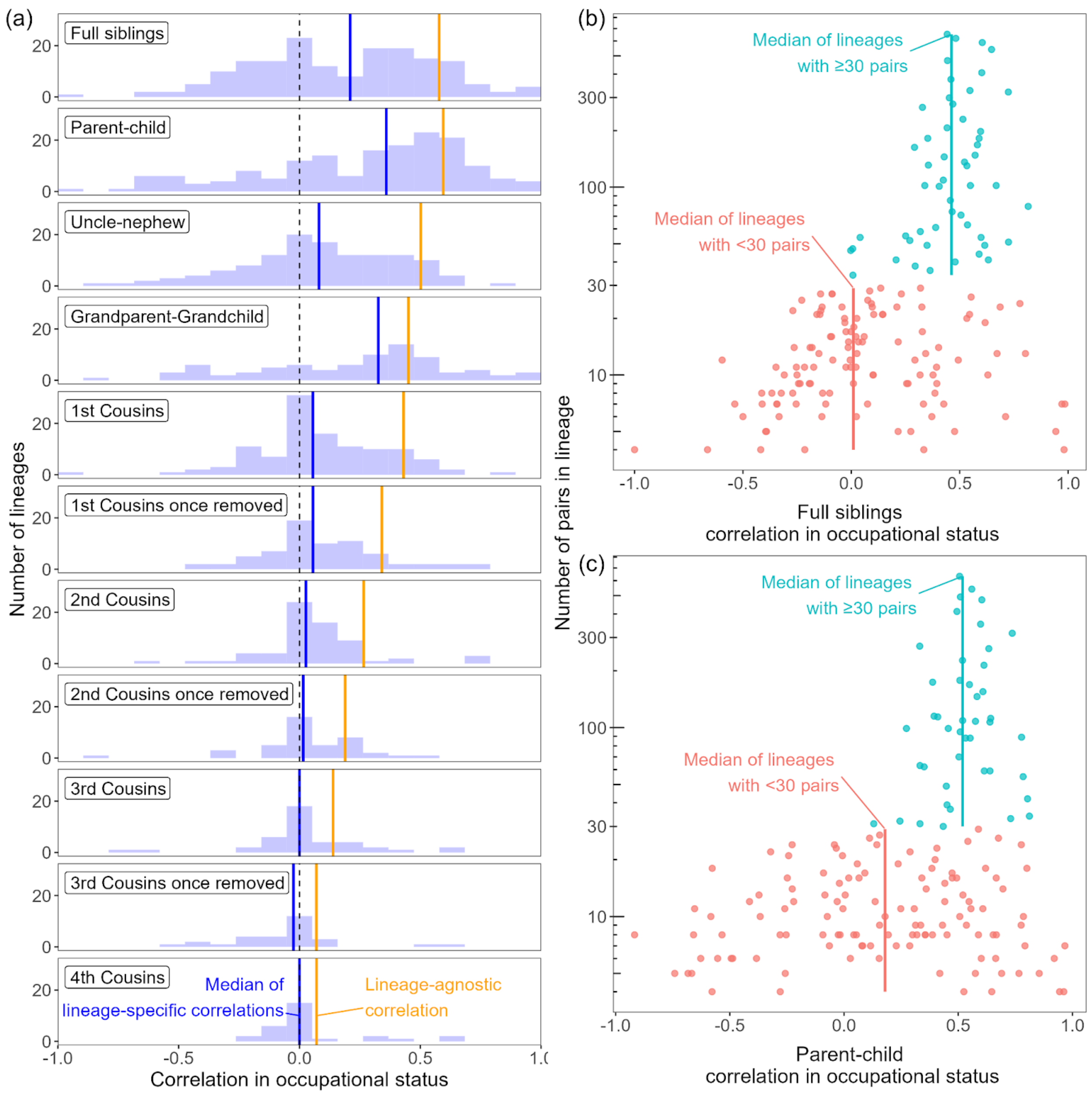


**Figure S2. Familial correlations in occupational status vary widely across surname lineages.**
**(a)** Comparing the “lineage-agnostic” correlations in occupational status (1780-1859) that (*1*) relies on (orange lines) with the distributions of lineage-specific correlations and their medians (blue lines). **(b,c)** Lineage-specific correlations in occupational status for first-degree relatives are associated with the representation of the lineage in (*1*). Lineages with ≥30 full sibling pairs **(b)** or father-son pairs **(c)** have a median trait correlation of 0.5, only slightly below the overall lineage-agnostic correlations shown in **(a)**. Lineages with <30 pairs in the data show much lower median correlations (~0.01 for full siblings, ~0.18 for father-son pairs).

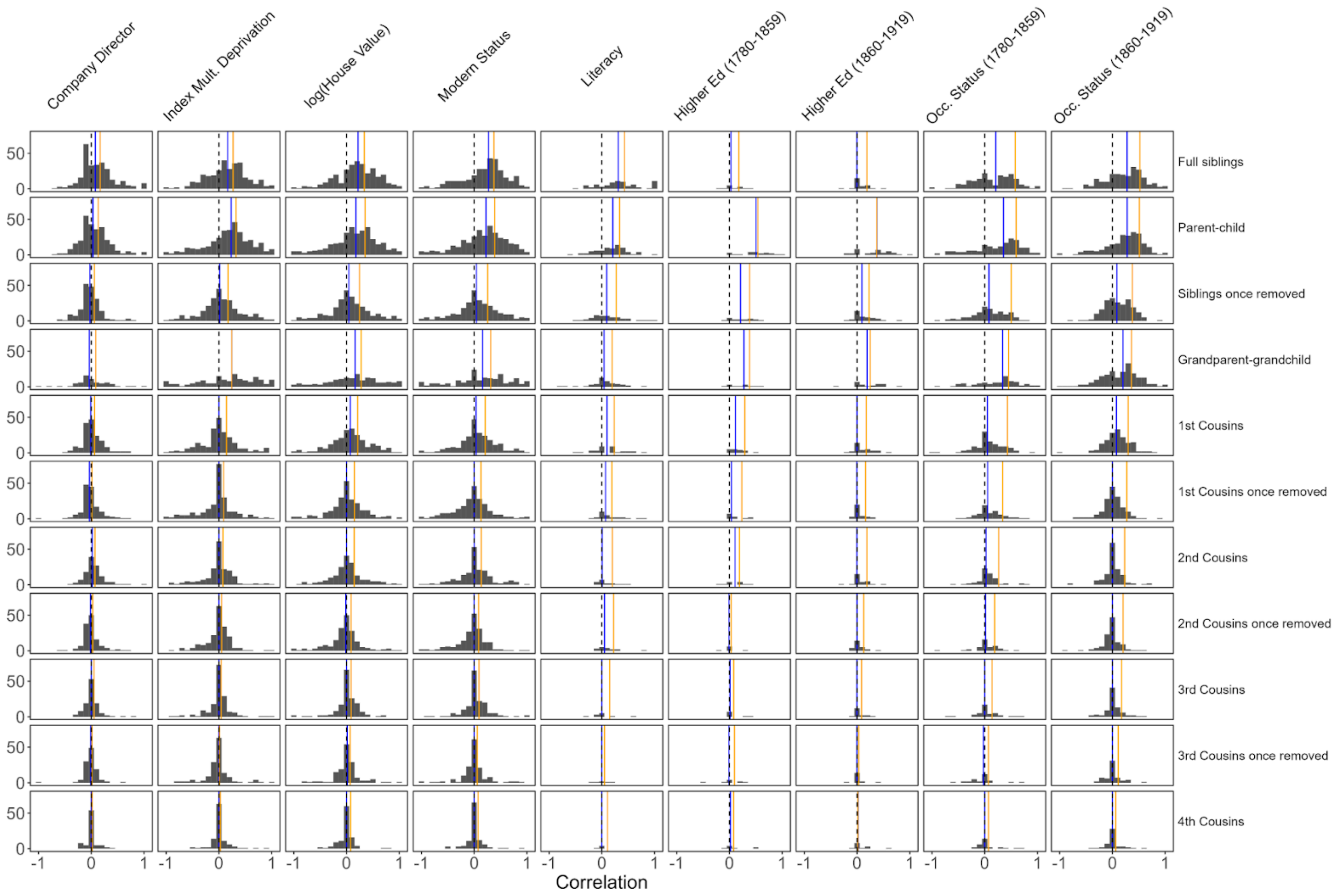


**Figure S3. Distributions of lineage-specific familial status correlations.**
Comparing the “lineage-agnostic” correlations presented in (*1*) (orange lines) with the distributions of lineage-specific correlations and their medians (blue lines), for measures of status analyzed in (*1*). Dashed black line marks zero. Many lineage-specific correlations for literacy and higher education could not be calculated (and are thus not plotted) due to there being no variation within the lineage. This is because most individuals did not go to university, and most individuals were literate.

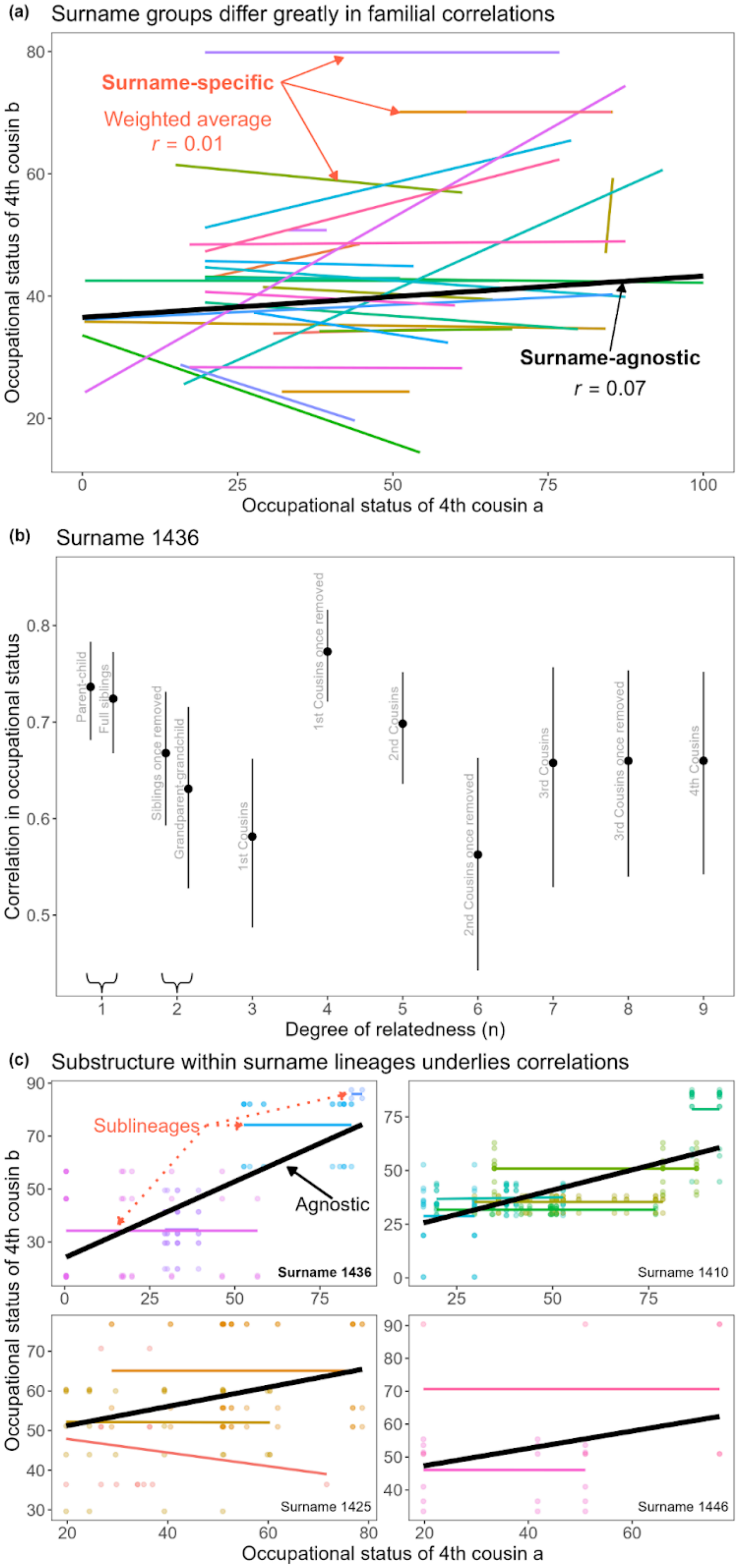


**Figure S4. Variation and substructure among and within surname lineages.** **(a)** When conditioning on surname lineage (indicated by colored lines), the association in occupational status between fourth cousins is highly variable. The weighted average of these correlations (where weights are determined by the number of pairs per lineage) is lower than the surname-agnostic correlation. **(b)** In one surname lineage (surname ID 1436), the surname-specific correlations are >0.5 for all relationship types, and there is no decay with genealogical distance, indicating that the model parameters estimated using lineage-agnostic correlation estimates cannot be considered representative of a given surname lineage. **(c)** Strong positive correlations within a surname lineage are attributable to substructure among those lineage members. Here we show fourth cousin data for four surname lineages. For each of these lineages, there are two or more sublineages (identified by their last common male ancestor) that differ in their average occupational status, but there is no within-sublineage correlation. All data shown in this figure are for individuals born 1780-1859.

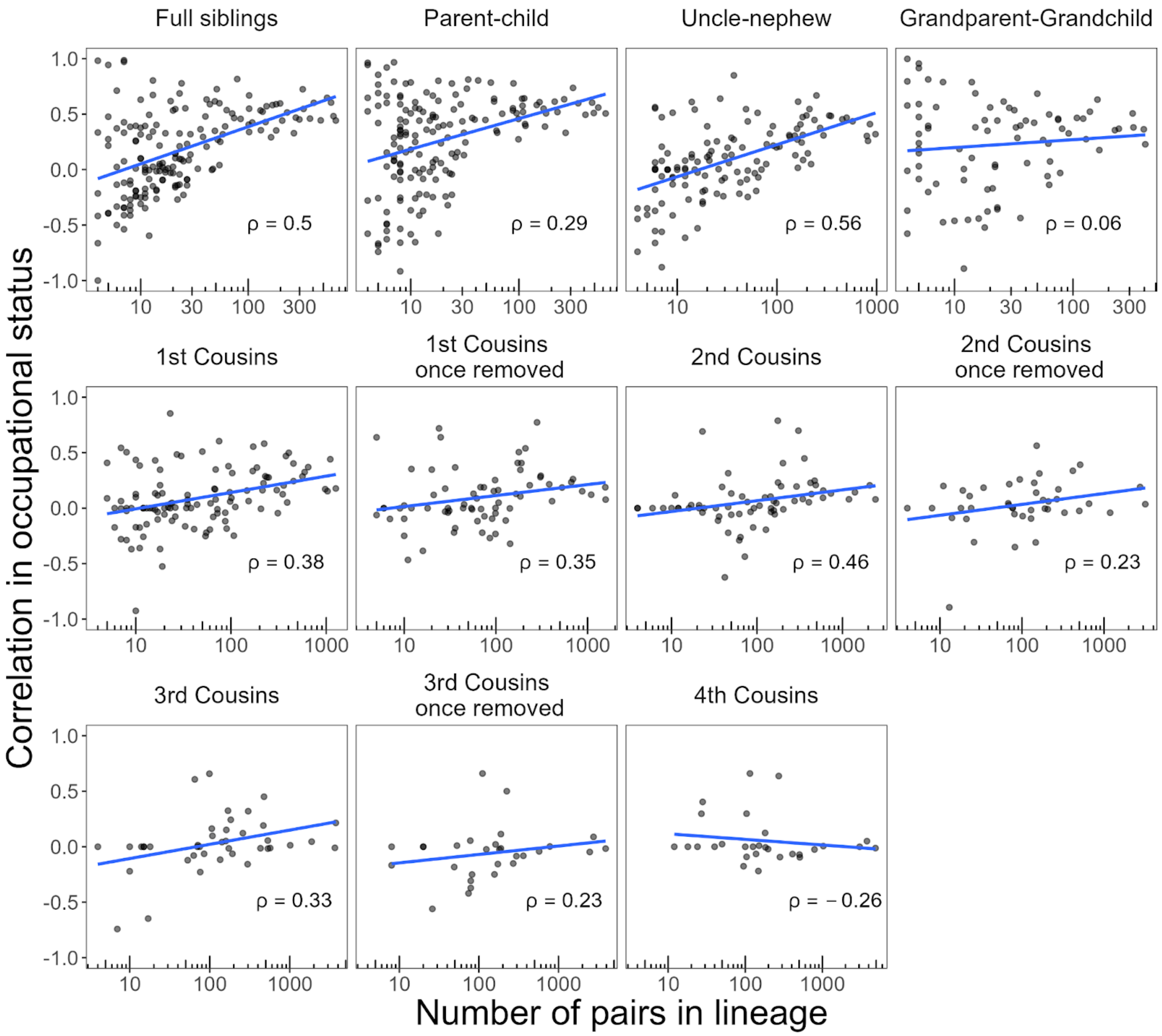


**Figure S5. Relationship between lineage-specific correlation in occupational status and lineage size, across relative types.**
The Spearman rank correlation (ρ) between these two variables is given in the bottom right corner of each panel. Blue lines show a linear fit to the data.

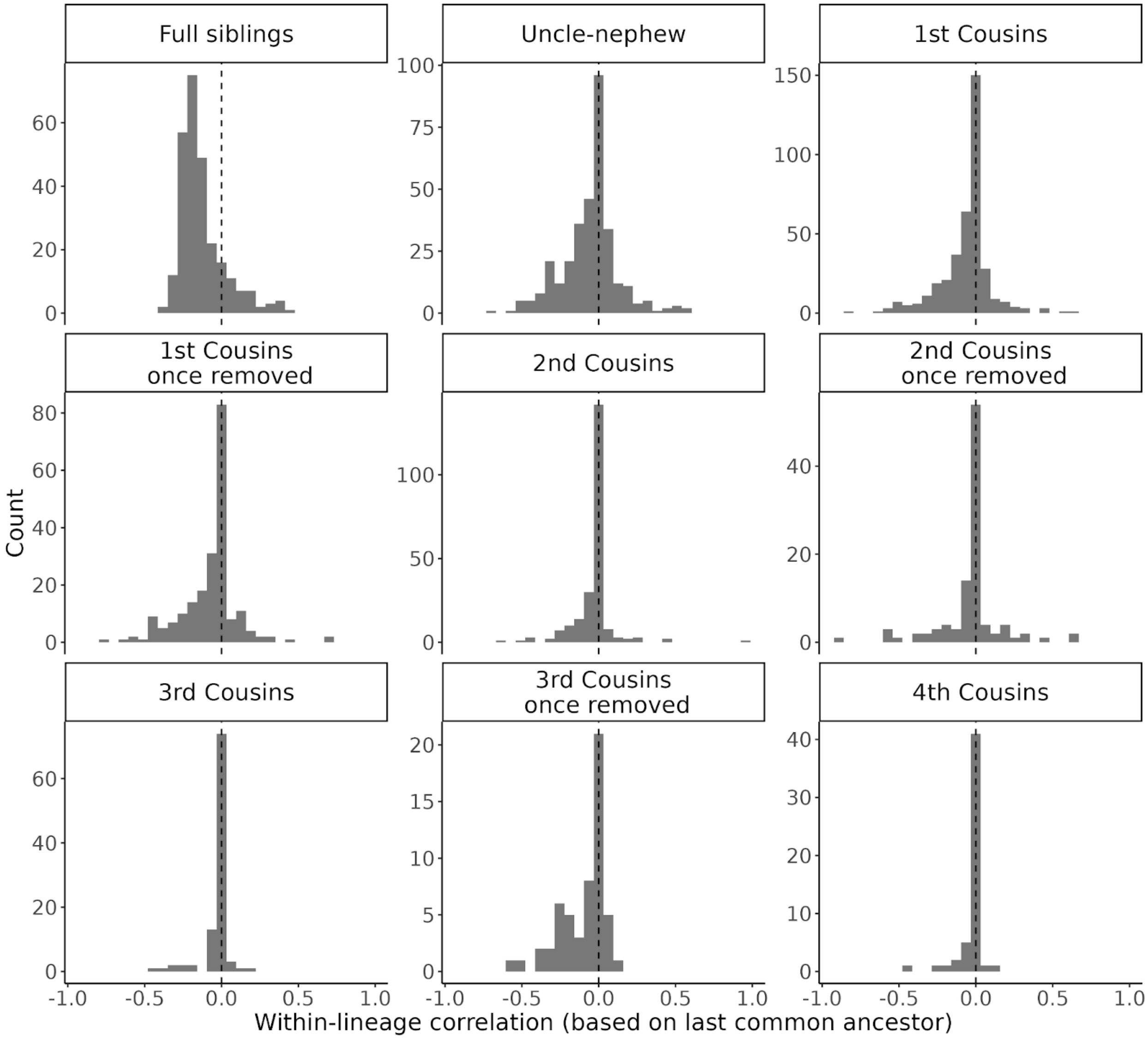


**Figure S6. Distributions of familial status correlations stratified by sublineage.** When conditioning on a single sublineage (defined by the last common ancestor of a set of relatives), the within-sublineage correlation in occupational status is typically near 0, for all genealogical relationships. Data are shown for sublineages in each relationship subset with at least 10 relative pairs. Note that vertical relationships are excluded because each relative pair includes an observation for the same father or grandfather, so there is no variance and thus a correlation cannot be calculated.

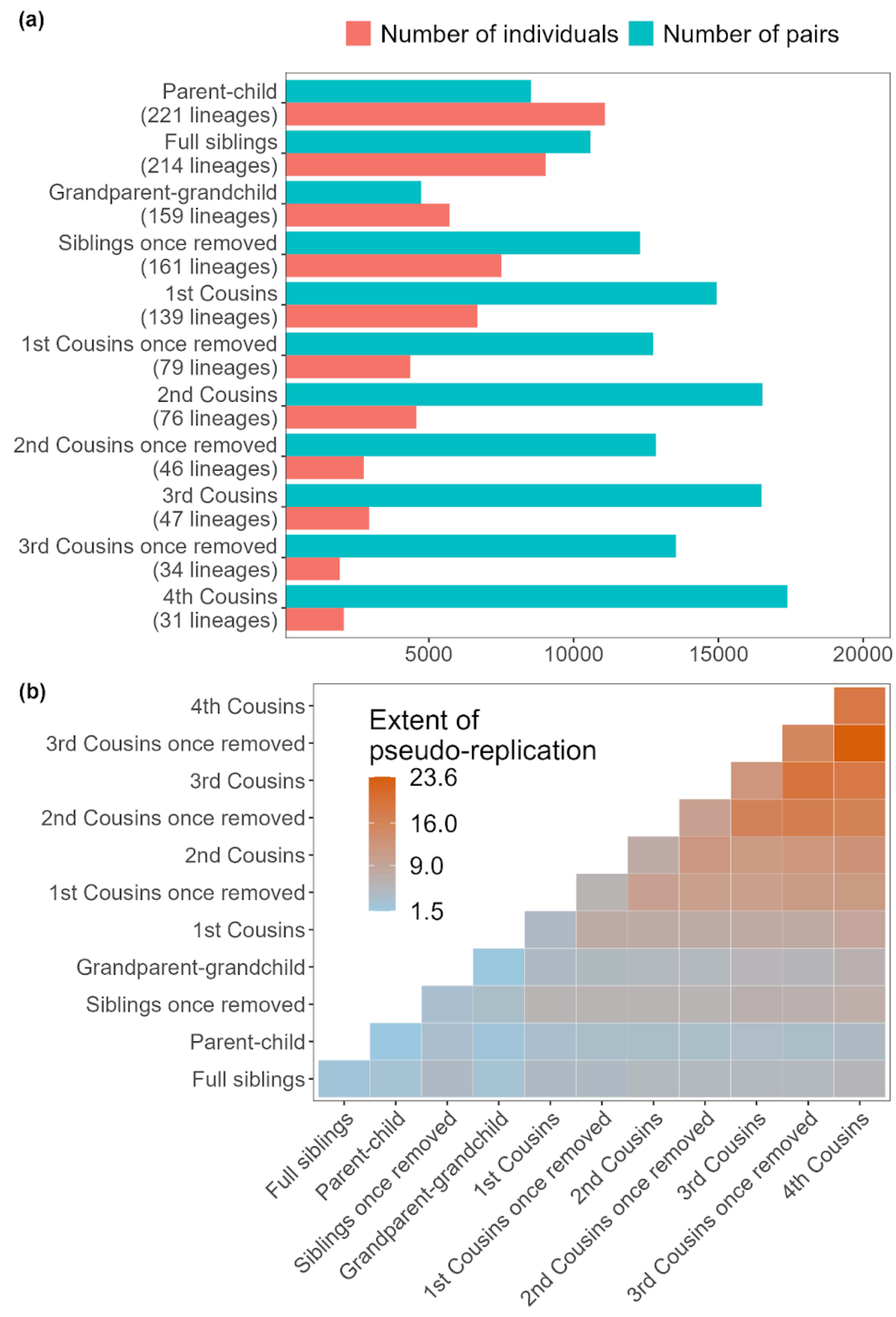


**Figure S7. Pseudoreplication of individuals in Clark (2023), illustrated here for the measure of occupational status (1780-1859).**
**(a)** the number of relative pairs increases with genealogical distance, even though the number of unique individuals and the number of surname lineages decreases, indicating increasing pseudoreplication among distant relatives. **(b)** We measure the "extent of pseudoreplication" as the average number of times an individual is included as a member of a pair in either the row category or the column category, across all individuals included in either of the two categories; this statistic ranges from 1.5 to 23.6 for these data. In the absence of pseudoreplication, this number would be 1.

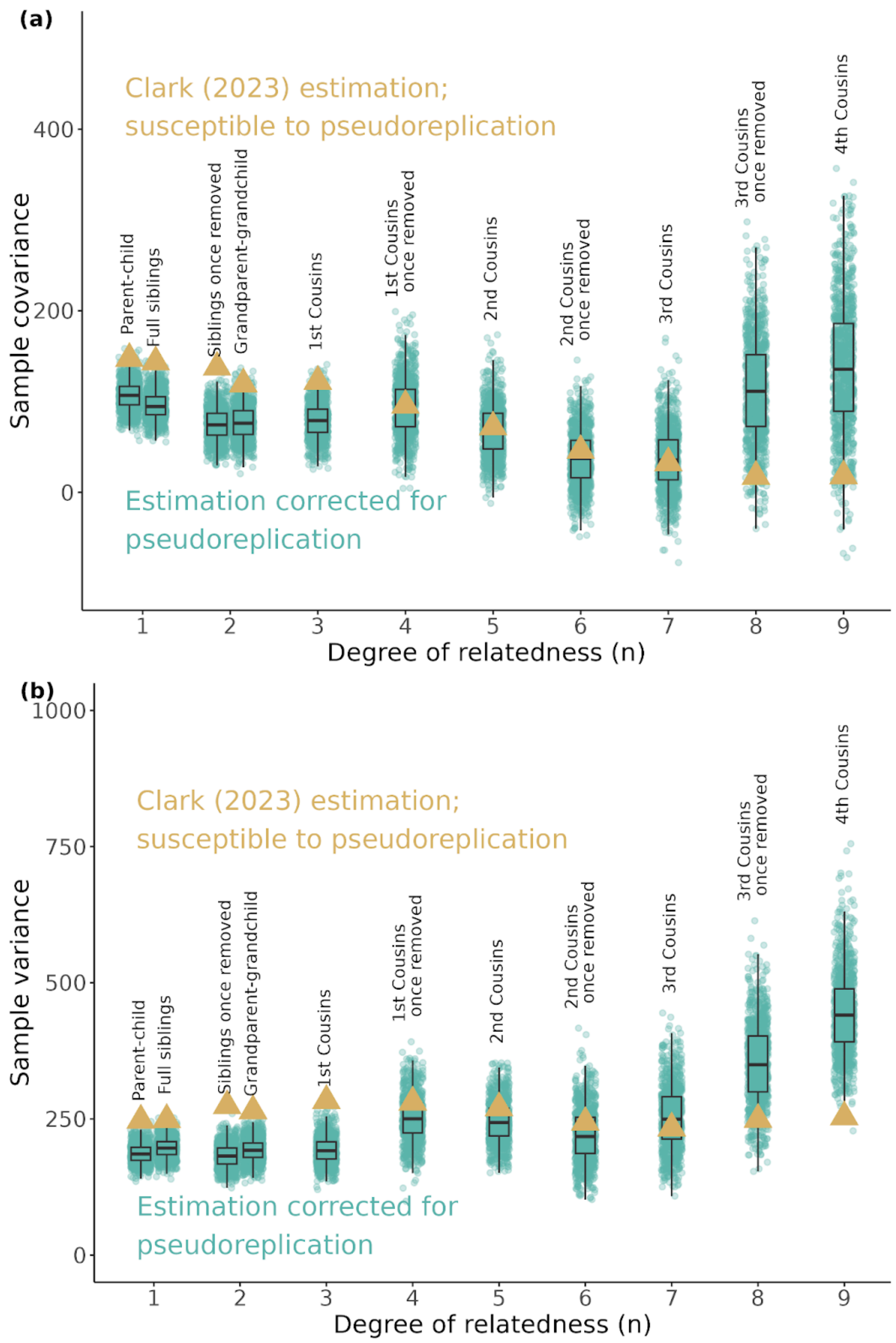


**Figure S8. Effect of pseudoreplication in Clark (2023) on variance and covariance estimates.**

Pseudoreplication alters both the covariance **(a)** and variance **(b)** of the data, which influences correlation estimates.

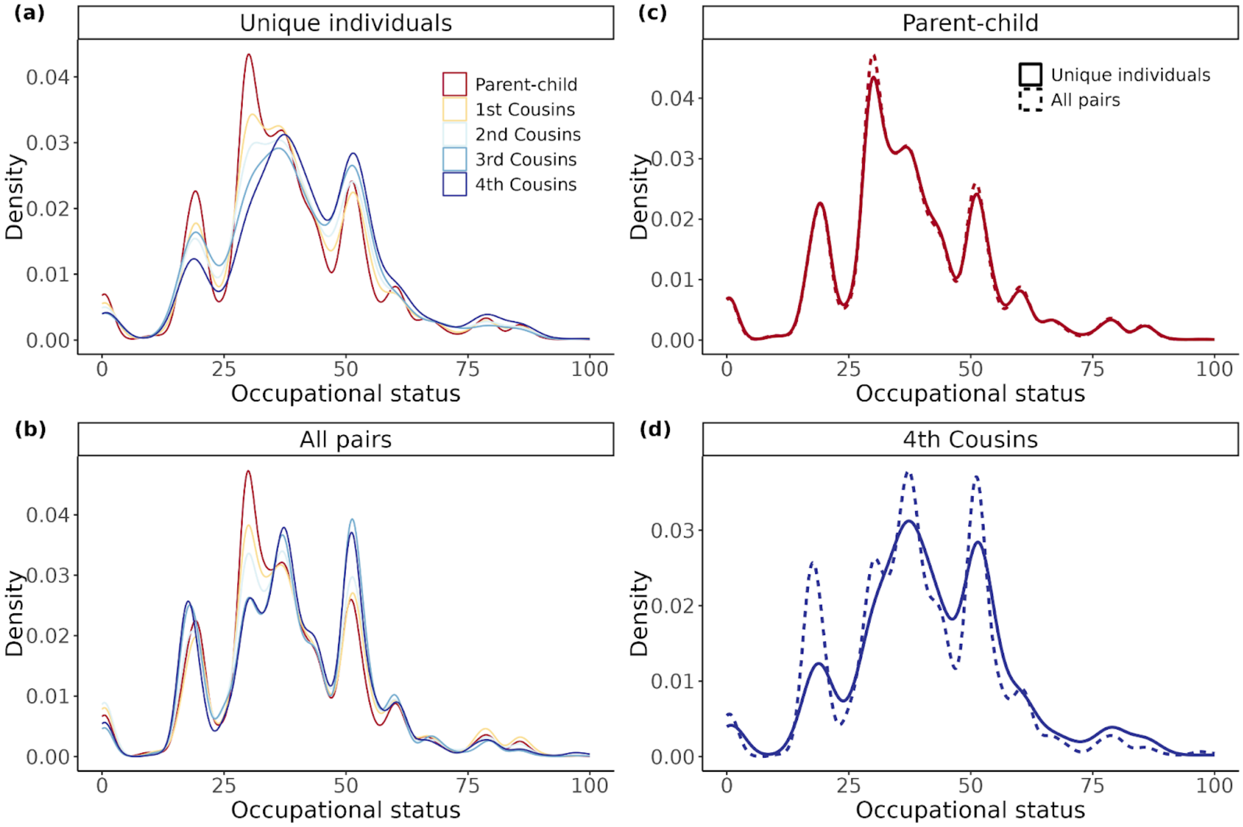


**Figure S9. Empirical distributions of occupational status scores vary with genealogical distance and are distorted by pseudoreplication.**

The method used to derive numerical scores for occupational status is described in **Supplementary Note 5**. Panel **(a)** shows the empirical distribution of occupational status scores considering only unique individuals in each relationship subset. Panel **(b)** shows the empirical distribution of these scores when including replicated records (as used in that status correlations reported in (*1*)). The distributions exhibit differences in abundance around different modal values (e.g. distant cousins tend to consist of relatively more individuals with occupational status scores near 50), and the overall mean and variance of these distributions tend to increase with genealogical distance, suggesting the different relative subsets are susceptible to sampling bias and/or come from different temporal populations (**Supplementary Note 6c**; **Fig. S10**). Direct comparisons of the distributions for unique individuals versus replicated records in father-son pairs **(c)** and fourth cousin pairs **(d)** show that pseudoreplication among distant relatives amplifies these distributional differences and affects the final estimates of familial correlations. All data shown are for men born 1780-1859.

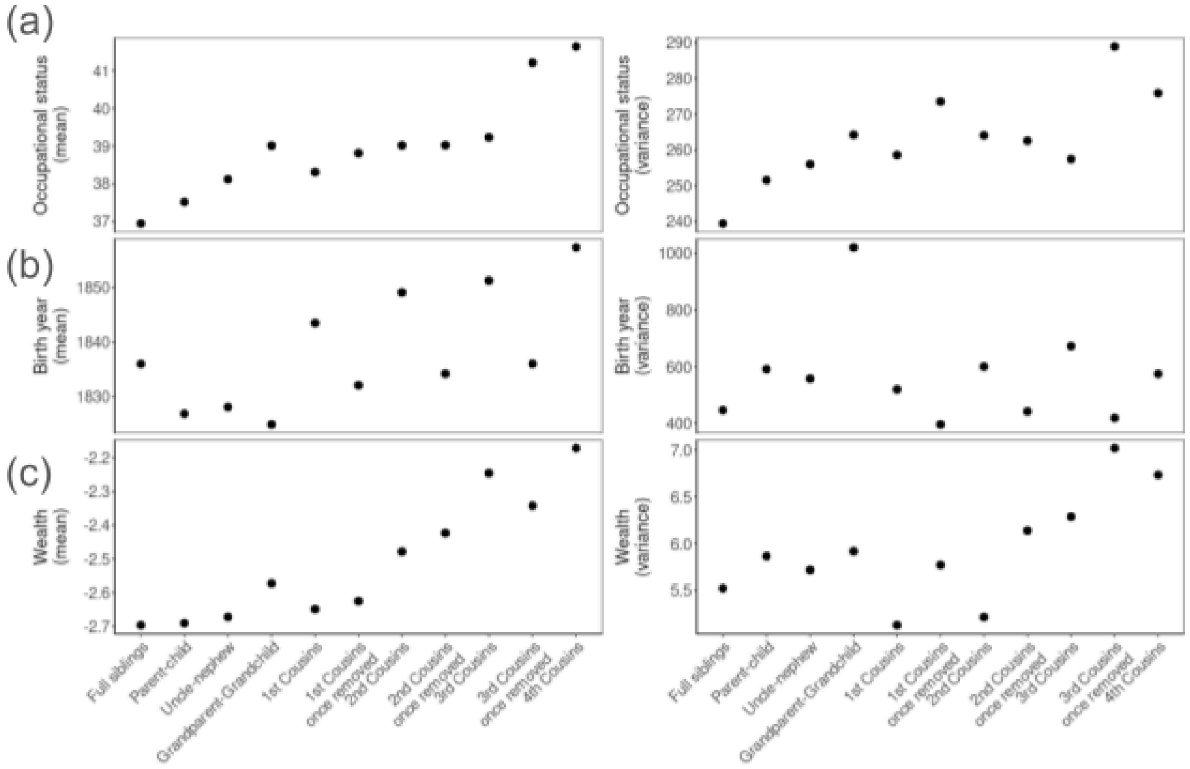


**Figure S10. Mean and variance of birth year and status measures across genealogical distance.**
**(a)** The mean and variance of occupational status (considering only values from unique individuals) increases with genealogical distance in the dataset of (*1*). Two potential explanations for these trends may be: **(b)** individuals represented in more distant relationship types in the data tend to have been born more recently so the trends are partly driven by unaccounted temporal changes in the population distribution of social status measures, or **(c)** biases in sampling or availability of genealogical data drive an enrichment of wealthier families among individuals represented in distant relative pairs.

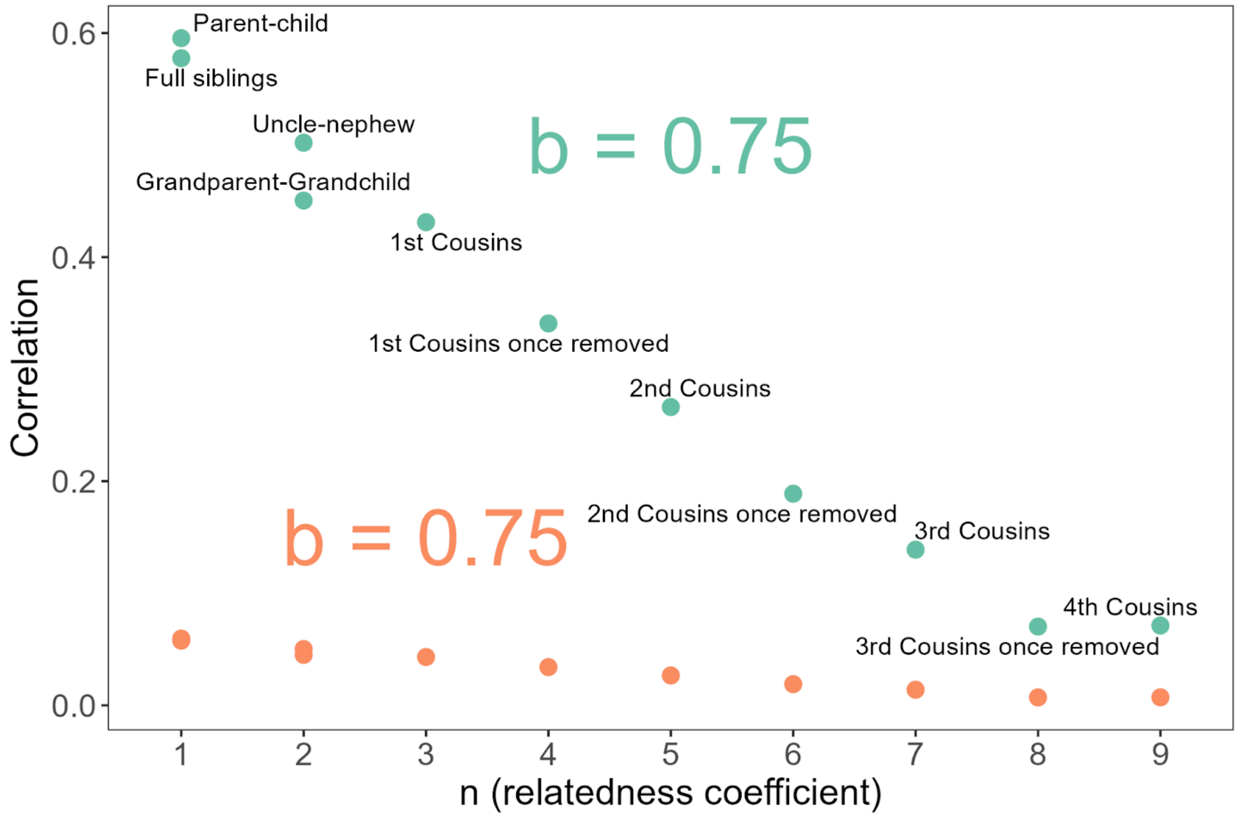


**Figure S11. The persistence parameter *b* is insensitive to the magnitude of familial correlations and describes only the *rate* at which they change as relatedness decreases.** As illustrated here with mock data, familial correlations in a status measure could increase or decrease by any degree with $b$ remaining unchanged (here, orange points are 10% of their corresponding points in green). For both, $R^2 = 0.96$.

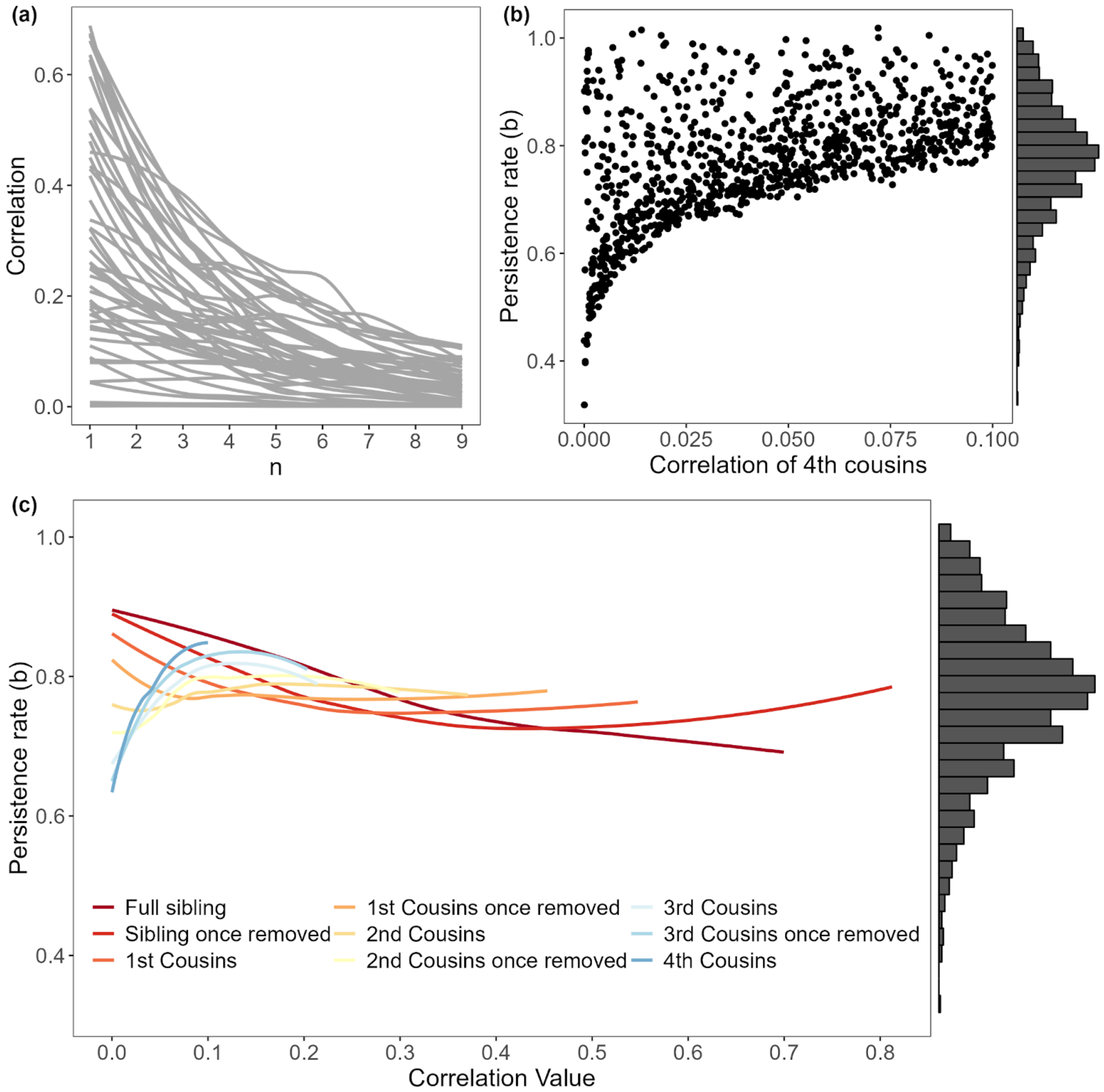


**Figure S12. Sensitivity of the persistence parameter, *b*, to changes in relative correlations.**

**(a)** Smoothed patterns of simulated familial correlations (only 50 out of 1000 simulations shown for clarity). **(b)** As the correlation of fourth cousins increases from zero, the minimum possible estimate of *b* increases sharply. **(c)** The sensitivity of the estimated *b* to different familial correlations. LOESS smoothed patterns of *estimated b ~ correlation value*, across all 1000 simulations, are plotted for each relationship. Small changes in the value of distant relative correlations result in relatively large changes in estimated *b*. See Supplementary Note 7 for simulation details.

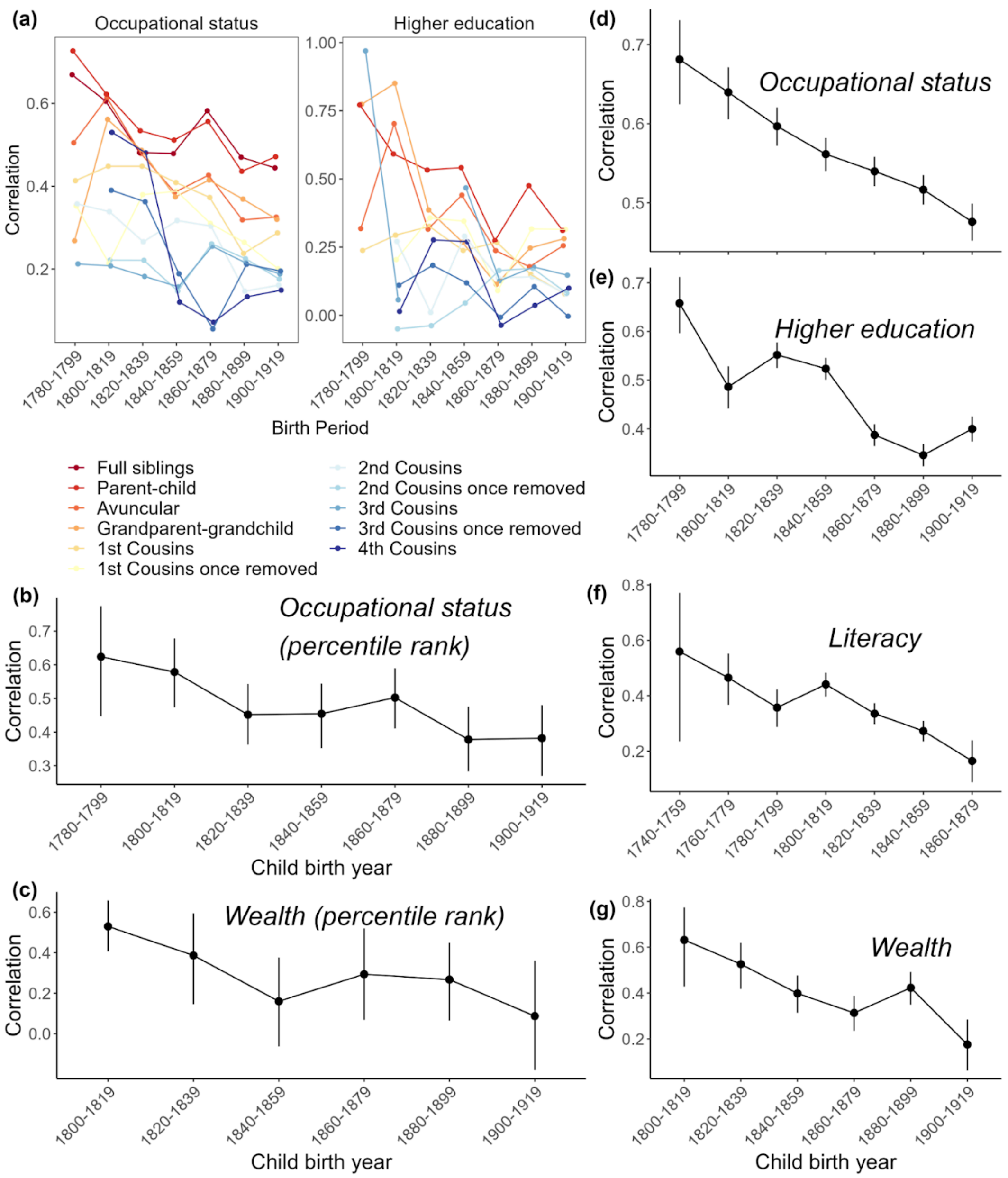


**Figure S13.Complementary analyses of temporal trends in familial correlations.** Compare to **Fig. 2c**. **(a)** Correlations (95% CI) in occupational status and higher education for all relationship types. Data were binned according to the birth year of the 2nd relative in each kin pair (for relationships that cross generations, this will be the younger generation). Higher education data for full siblings is not plotted, as correlations calculated from the raw data do not match those published in (*1*); we were not able to obtain corrected data from the author at time of this publication. **(b-c)** Change in father-son correlations in occupational status (b) and wealth (c) across 1780-1919, using percentile ranks of each individual within each time period. As in the main analysis, to mitigate pseudoreplication, we calculated correlations in (a-c) using one pair from each surname. Shown are mean correlations across 500 bootstrap iterations of correlation estimation. **(d-g)** Parent-offspring correlations (95% CI) over time, calculated from raw data and not corrected for pseudoreplication. (d) Occupational status 1780-1919; (e) higher education 1780-1919; (f) literacy 1740-1879; (g) wealth 1800-1919.

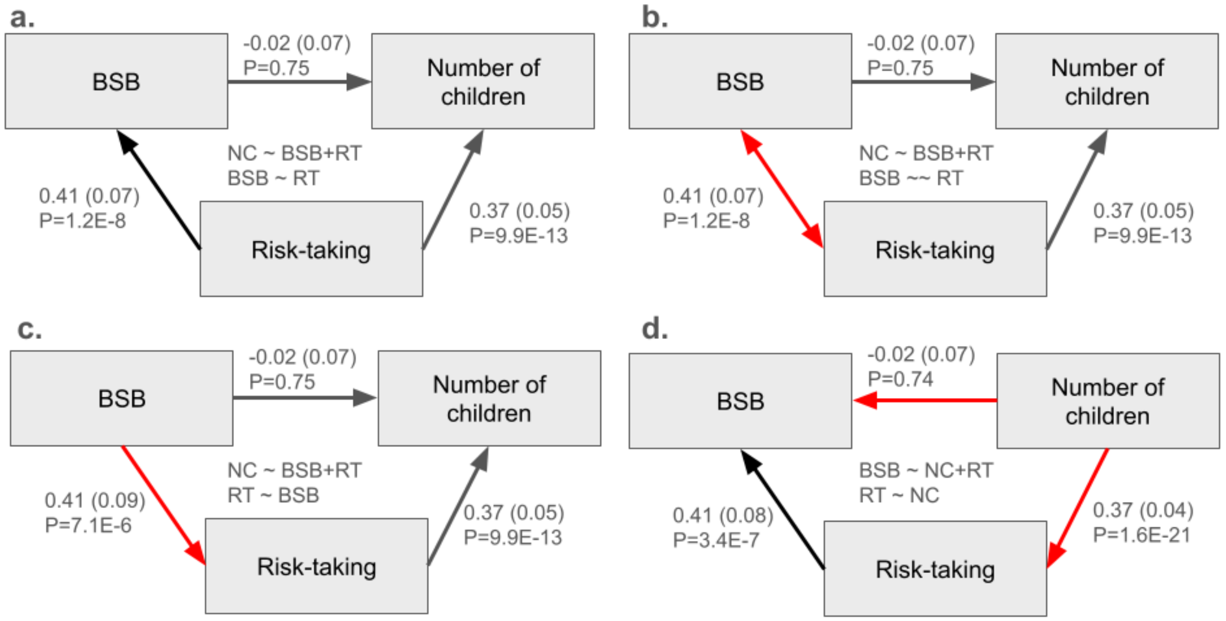


**Figure S14. Path diagrams depicting results of Genomic SEM models with different postulated causal relationships between BSB, risk-taking, and number of children.** Partial genetic correlations (with SE in parentheses) and their p-values are shown near each edge, and the model syntax is given at the center of the diagram. (a) Path diagram for our reproduction of the Genomic SEM model postulated in (29). (b-d) Path diagrams depicting alternative postulated causal relationships between BSB, risk-taking, and number of children. Red arrows indicate hypothesized causal paths that differ from the model shown in (a). (b) path diagram where we model the genetic covariance between BSB and risk-taking (i.e. explicitly modeling horizontal pleiotropy). (c) Path diagram for a model where the causal path between risk-taking and BSB has been reversed from that of (a). (d) Path diagram where we reverse the causal paths pointing from BSB and risk-taking to the number of children as in (a), but do not change the path from risk-taking to BSB. In all four models, the point estimates for the partial correlations are identical, differing only slightly in standard errors and p-values. This comparison demonstrates that the results presented in (29) cannot be interpreted as support of any particular mode of genetic causation.

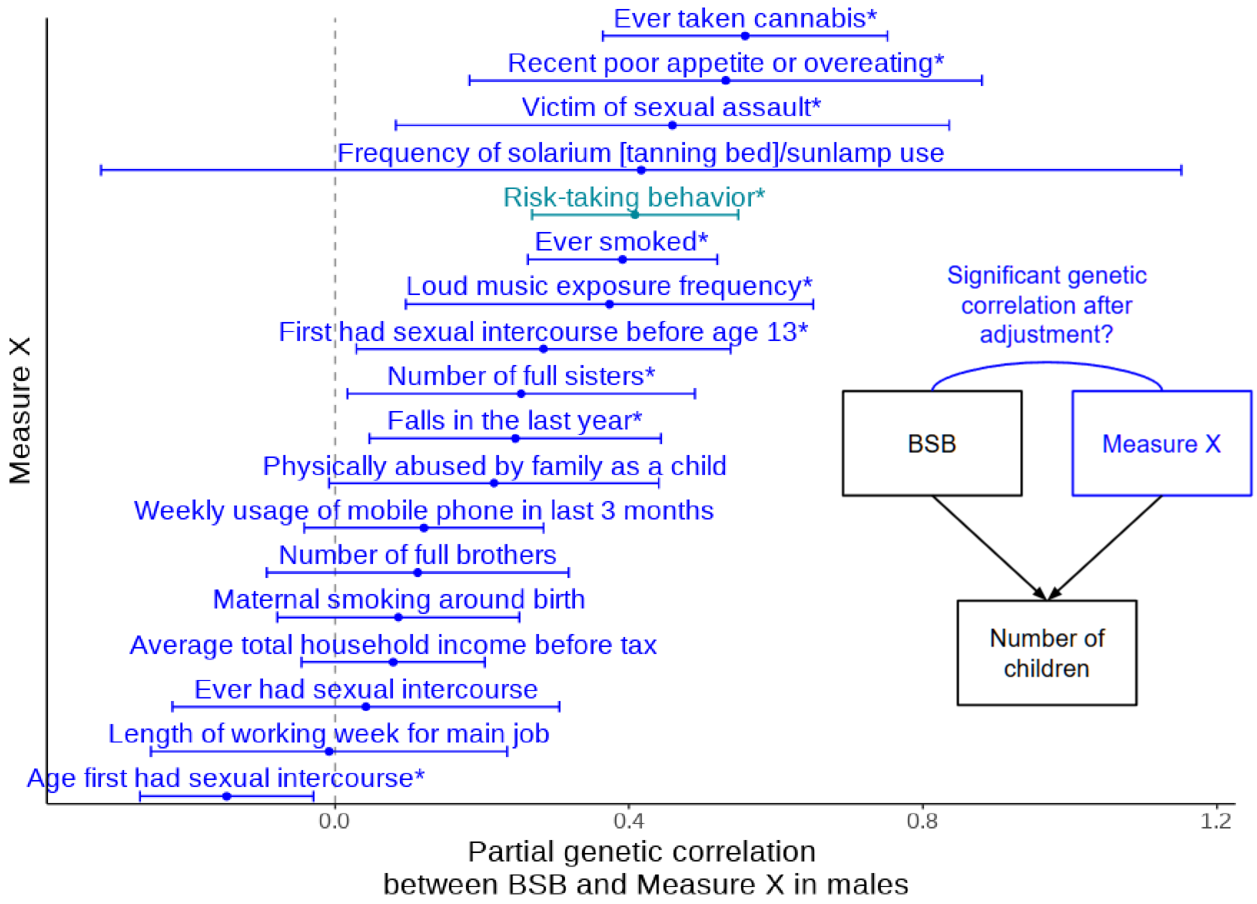


**Figure S15. Partial genetic correlations between BSB and the measures considered in the alternative Genomic SEM models described in Fig. 3b.**

Whereas Fig. 3b shows the partial genetic correlations between BSB and number of children (after adjusting for each measure), here we show the partial correlations between BSB and each of those measures (corresponding to the blue path in the inset diagram). Error bars indicate 95% confidence intervals on the estimated partial genetic correlations. Measures that have significant partial genetic correlations with BSB (unadjusted Genomic SEM P-value $< 0.05$) are denoted with an asterisk.

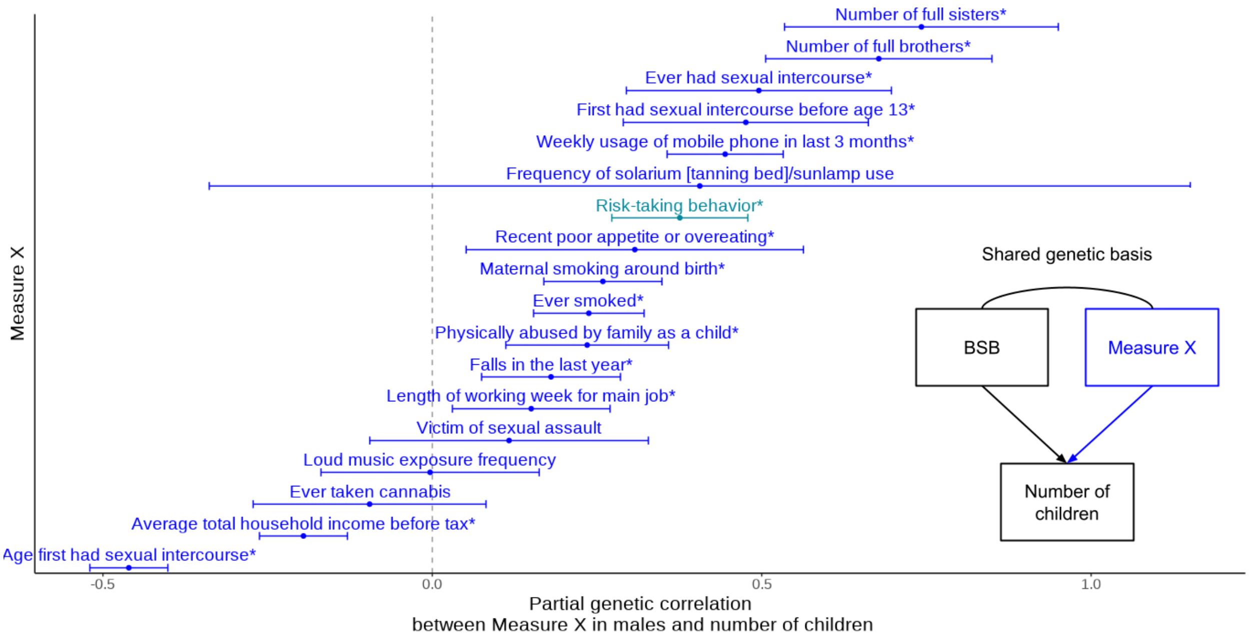


**Figure S16. Partial genetic correlations between number of children and the measures considered in the alternative Genomic SEM models described in Fig. 3b.**
Whereas Fig. 3b shows the partial genetic correlations between BSB and number of children (after adjusting for each measure), here we show the partial correlations between number of children and each of those measures (corresponding to the blue path in the inset diagram). Error bars indicate 95% confidence intervals on the estimated partial genetic correlations. Measures that have significant partial genetic correlations with number of children (unadjusted Genomic SEM P-value < 0.05) are denoted with an asterisk.

### References


1. G. Clark, The inheritance of social status: England, 1600 to 2022. *Proc. Natl. Acad. Sci. U. S. A.* **120**, e2300926120 (2023).

2. A. Gimelfarb, A general linear model for the genotypic covariance between relatives under assortative mating. *J. Math. Biol.* **13**, 209–226 (1981).

3. R. A. Fisher, The Correlation between Relatives on the Supposition of Mendelian Inheritance. *Trans. R. Soc. Edinb.* **52**, 399–433 (1918).

4. C. R. Cloninger, J. Rice, T. Reich, Multifactorial inheritance with cultural transmission and assortative mating. II. a general model of combined polygenic and cultural inheritance. *Am. J. Hum. Genet.* **31**, 176–198 (1979).

5. M. D. Collado, I. Ortuño-Ortín, J. Stuhler, Estimating Intergenerational and Assortative Processes in Extended Family Data. *Rev. Econ. Stud.* **90**, 1195–1227 (2023).

6. B. W. Domingue, J. Fletcher, D. Conley, J. D. Boardman, Genetic and educational assortative mating among US adults. *Proc. Natl. Acad. Sci. U. S. A.* **111**, 7996–8000 (2014).

7. E. T. Akimova, T. Wolfram, X. Ding, F. C. Tropf, M. C. Mills, Polygenic predictions of occupational status GWAS elucidate genetic and environmental interplay for intergenerational status transmission, careers, and health, *bioRxiv* (2023)p. 2023.03.31.534944.

8. L. Yengo, M. R. Robinson, M. C. Keller, K. E. Kemper, Y. Yang, M. Trzaskowski, J. Gratten, P. Turley, D. Cesarini, D. J. Benjamin, N. R. Wray, M. E. Goddard, J. Yang, P. M. Visscher, Imprint of assortative mating on the human genome. *Nat Hum Behav* **2**, 948–954 (2018).

9. M. R. Robinson, A. Kleinman, M. Graff, A. A. E. Vinkhuyzen, D. Couper, M. B. Miller, W. J. Peyrot, A. Abdellaoui, B. P. Zietsch, I. M. Nolte, J. V. van Vliet-Ostaptchouk, H. Snieder, S. E. Medland, N. G. Martin, P. K. E. Magnusson, W. G. Iacono, M. McGue, K. E. North, J. Yang, P. M. Visscher, Genetic evidence of assortative mating in humans. *Nature Human Behaviour* **1**, 1–13 (2017).

10. A. Okbay, Y. Wu, N. Wang, H. Jayashankar, M. Bennett, S. M. Nehzati, J. Sidorenko, H. Kweon, G. Goldman, T. Gjorgjieva, Y. Jiang, B. Hicks, C. Tian, D. A. Hinds, R. Ahlskog, P. K. E. Magnusson, S. Oskarsson, C. Hayward, A. Campbell, D. J. Porteous, J. Freese, P. Herd, 23andMe Research Team, Social Science Genetic Association Consortium, C. Watson, J. Jala, D. Conley, P. D. Koellinger, M. Johannesson, D. Laibson, M. N. Meyer, J. J. Lee, A. Kong, L. Yengo, D. Cesarini, P. Turley, P. M. Visscher, J. P. Beauchamp, D. J. Benjamin, A. I. Young, Polygenic prediction of educational attainment within and between families from genome-wide association analyses in 3 million individuals. *Nat. Genet.* **54**, 437–449 (2022).

11. G. Clark, N. Cummins, M. Curtis, The Mismeasure of Man: Why Intergenerational Occupational Mobility is Much Lower than Conventionally Measured, England, 1800-2021 (2022). https://papers.ssrn.com/abstract=4144664.

12. P. S. Lambert, R. L. Zijdeman, M. H. D. Van Leeuwen, I. Maas, K. Prandy, The Construction

of HISCAM: A Stratification Scale Based on Social Interactions for Historical Comparative Research. *Historical Methods: A Journal of Quantitative and Interdisciplinary History* **46**, 77–89 (2013).

13. L. A. Goodman, Multiplicative Models for the Analysis of Occupational Mobility Tables and Other Kinds of Cross-Classification Tables. *Am. J. Sociol.* **84**, 804–819 (1979).

14. M. S. Lewis-Beck, A. Bryman, T. F. Liao, *The SAGE Encyclopedia of Social Science Research Methods* (Sage Publications, Inc., 2011).

15. S. H. Hurlbert, Pseudoreplication and the Design of Ecological Field Experiments. *Ecol. Monogr.* **54**, 187–211 (1984).

16. S. Scarr-Salapatek, Race, Social Class, and IQ. *Science* **174**, 1285–1295 (1971).

17. D. C. Rowe, K. C. Jacobson, E. J. C. G. Van den Oord, Genetic and environmental influences on vocabulary IQ: Parental education level as moderator. *Child Dev.* **70**, 1151–1162 (1999).

18. E. Turkheimer, K. P. Harden, B. d'Onofrio, I. I. Gottesman, The Scarr--Rowe interaction between measured socioeconomic status and the heritability of cognitive ability. *Experience and development: A festschrift in honor of Sandra Wood Scarr* **2**, 81–97 (2009).

19. E. Turkheimer, E. E. Horn, "Interactions Between Socioeconomic Status and Components of Variation in Cognitive Ability" in *Behavior Genetics of Cognition Across the Lifespan*, D. Finkel, C. A. Reynolds, Eds. (Springer New York, New York, NY, 2014), pp. 41–68.

20. E. M. Tucker-Drob, T. C. Bates, Large Cross-National Differences in Gene × Socioeconomic Status Interaction on Intelligence. *Psychol. Sci.* **27**, 138–149 (2016).

21. E. J. Giangrande, E. Turkheimer, Race, Ethnicity, and the Scarr-Rowe Hypothesis: A Cautionary Example of Fringe Science Entering the Mainstream. *Perspect. Psychol. Sci.* **17**, 696–710 (2022).

22. M. W. Feldman, F. B. Christiansen, S. P. Otto, Gene-culture co-evolution: teaching, learning, and correlations between relatives. *Isr. J. Ecol. Evol.* **59**, 72–91 (2013).

23. O. Kolodny, M. W. Feldman, A. Lotem, Y. Ram, Differential application of cultural practices at the family and individual levels may alter heritability estimates. *Behav. Brain Sci.* **45**, e167 (2022).

24. S. E. Lazic, The problem of pseudoreplication in neuroscientific studies: is it affecting your analysis? *BMC Neurosci.* **11**, 5 (2010).

25. N. A. Rosenberg, J. M. Vanliere, Replication of genetic associations as pseudoreplication due to shared genealogy. *Genet. Epidemiol.* **33**, 479–487 (2009).

26. O. Causa, Å. Johansson, "Intergenerational Social Mobility" (Organisation for Economic Co-Operation and Development (OECD), 2009); https://doi.org/10.1787/223106258208.

27. R. Chetty, N. Hendren, P. Kline, E. Saez, N. Turner, Is the United States Still a Land of Opportunity? Recent Trends in Intergenerational Mobility. *Am. Econ. Rev.* **104**, 141–147 (2014).

28. P. A. Longley, J. van Dijk, T. Lan, The geography of intergenerational social mobility in Britain. *Nat. Commun.* **12**, 6050 (2021).

29. S. Song, J. Zhang, Genetic variants underlying human bisexual behavior are reproductively advantageous. *Sci Adv* **10**, eadj6958 (2024).

30. C. Bycroft, C. Freeman, D. Petkova, G. Band, L. T. Elliott, K. Sharp, A. Motyer, D. Vukcevic, O. Delaneau, J. O'Connell, A. Cortes, S. Welsh, A. Young, M. Effingham, G. McVean, S. Leslie, N. Allen, P. Donnelly, J. Marchini, The UK Biobank resource with deep phenotyping and genomic data. *Nature* **562**, 203–209 (2018).

31. A. D. Grotzinger, M. Rhemtulla, R. de Vlaming, S. J. Ritchie, T. T. Mallard, W. D. Hill, H. F. Ip, R. E. Marioni, A. M. McIntosh, I. J. Deary, P. D. Koellinger, K. P. Harden, M. G. Nivard, E. M. Tucker-Drob, Genomic structural equation modelling provides insights into the multivariate genetic architecture of complex traits. *Nat Hum Behav* **3**, 513–525 (2019).

32. C. C. Chang, C. C. Chow, L. C. Tellier, S. Vattikuti, S. M. Purcell, J. J. Lee, Second-generation PLINK: rising to the challenge of larger and richer datasets. *Gigascience* **4**, 7 (2015).

33. J. Mbatchou, L. Barnard, J. Backman, A. Marcketta, J. A. Kosmicki, A. Ziyatdinov, C. Benner, C. O'Dushlaine, M. Barber, B. Boutkov, L. Habegger, M. Ferreira, A. Baras, J. Reid, G. Abecasis, E. Maxwell, J. Marchini, Computationally efficient whole-genome regression for quantitative and binary traits. *Nat. Genet.* **53**, 1097–1103 (2021).

34. B. Bulik-Sullivan, H. K. Finucane, V. Anttila, A. Gusev, F. R. Day, P.-R. Loh, ReproGen Consortium, Psychiatric Genomics Consortium, Genetic Consortium for Anorexia Nervosa of the Wellcome Trust Case Control Consortium 3, L. Duncan, J. R. B. Perry, N. Patterson, E. B. Robinson, M. J. Daly, A. L. Price, B. M. Neale, An atlas of genetic correlations across human diseases and traits. *Nat. Genet.* **47**, 1236–1241 (2015).
